# p53 restoration in small cell lung cancer identifies a latent Cyclophilin-dependent necrosis mechanism

**DOI:** 10.1101/2022.07.22.501202

**Authors:** Jonuelle Acosta, Qinglan Li, Nelson F. Freeburg, Nivitha Murali, Grant P. Grothusen, Michelle Cicchini, Hung Mai, Amy C. Gladstein, Keren M. Adler, Katherine R. Doerig, Jinyang Li, Miguel Ruiz-Torres, Kimberly L. Manning, Ben Z. Stanger, Luca Busino, Liling Wan, David M. Feldser

## Abstract

The p53 tumor suppressor regulates multiple context-dependent tumor suppressive programs. Although p53 is mutated in ∼90% of small cell lung cancer (SCLC) tumors, how p53 mediates tumor suppression in this context is unknown. Here, using a mouse model of SCLC in which endogenous p53 expression can be conditionally and temporally regulated, we show that SCLC tumors maintain a requirement for p53 inactivation. However, we identified tumor subtype heterogeneity between SCLC tumors such that p53 reactivation induces a canonical senescence response in a subset of tumors, while, in others, p53 induces a non-apoptotic form of cell death that culminates in necrosis. We pinpointed the cyclophilin family of peptidyl prolyl *cis*-*trans* isomerases as critical determinants of a p53-induced transcriptional program that is specific to SCLC tumors and cell lines that are poised to undergo p53-mediated necrosis. Importantly, inhibition of cyclophilin isomerase activity suppresses SCLC subtype-specific p53-mediated death by limiting p53 transcriptional output without impacting chromatin binding. Our study demonstrates that intertumoral heterogeneity in SCLC can influence the biological response to p53 restoration, describes a novel mechanism of p53-regulated necrotic cell death, and uncovers new targets for the treatment of this most-recalcitrant tumor type.

## Introduction

Small cell lung cancer (SCLC) is the most lethal subtype of lung cancer and comprises ∼15% of all lung cancer cases [1, 2]. The long-standing standard of care treatment for SCLC has been platinum-based chemotherapy, which leads to significant tumor shrinkage in up to 80% of patients [3]. However, most patients later present with a recurrence of disease and succumb to treatment-resistant tumors and metastases. Recent studies have unveiled that SCLC is not a homogeneous disease and is instead comprised of four distinct molecular subtypes based on expression of lineage-defining transcription factors: *ASCL1* (SCLC-A), *NEUROD1* (SCLC-N), *POU2F3* (SCLC-P), or *YAP1* (SCLC-Y) [4]. These molecular subtypes have also been associated with distinct subtype-specific therapeutic vulnerabilities that can be used in combination with traditional chemotherapeutic treatment to prevent disease recurrence and enhance survival [5–9]. Cell types of origin influence tumor evolution and metastatic progression, further contributing to intertumoral SCLC heterogeneity [10]. These insights highlight the need to better understand how SCLC heterogeneity influences etiology of the disease, which includes the near-universal persistent selective requirement for key cancer driving mutations [11].

Together with the retinoblastoma (Rb) tumor suppressor, p53 is inactivated in the vast majority (>90%) of SCLC cases [11]. p53 is a pleiotropic tumor suppressor gene that responds to a wide array of genotoxic and cellular stresses to effectuate diverse tumor suppressive programs such as activation of apoptosis and senescence, as well as metabolic programs that may be essential for tumor suppression [12, 13]. Identifying the contexts in which p53 induces one tumor suppressive program over another remains a challenge but may hold the key to establishing novel, contextual, therapeutic strategies. Mouse models of cancer that allow for the conditional restoration of p53 function after tumor formation have identified tissue-dependent contexts where p53 preferentially activates cytostatic or cytotoxic programs [14]. In spontaneous models of B or T-cell lymphoma, p53 restoration induces canonical caspase-dependent apoptosis and rapid tumor regression [15, 16]. In contrast, restoration of p53 in soft tissue sarcomas and hepatocellular carcinoma induces senescence which is followed by immune-mediated tumor clearance [16, 17]. Similarly, p53 restoration in a model of Kras-driven lung adenocarcinoma also induces a senescence response followed by immune-mediated clearance, but only in advanced tumor lesions [18–20]. These studies demonstrate that p53 controls distinct tumor suppressive programs in specific cancer types, highlight the need to better understand how p53 mediates tumor suppression in distinct contexts, and illustrate the power of gene-reactivation strategies to elucidate latent mechanisms of tumor suppression.

Here we uncover multiple cancer cell autonomous roles of p53 that are selectively-activated in distinct SCLCs. We demonstrate that p53 can drive a canonical senescence program in a subset of SCLC while inducing a non-canonical form of necrotic cell death in others. This cell death program is associated with a distinct transcriptional output after p53 reactivation that was unexpectedly dependent on multiple members of the cyclophilin family of peptidyl proline isomerases. Importantly, without affecting DNA binding directly, cyclophilin inhibition selectively limited p53-target gene expression specifically in SCLCs that are fated to undergo necrotic cell death after p53 restoration. Additionally, we present evidence that distinct SCLC tumor subtypes may underlie the divergent cell fates that occur after p53 restoration.

## Results

### p53 restoration limits growth and metastatic spread of autochthonous SCLC

To gain insight into the requirement of sustained p53 inactivation in SCLC, we employed a *Trp53*^XTR^ allele that we developed to turn off p53 expression during tumor development and then restore p53 expression in established, autochthonous cancers in the mouse [21]. *Rb1*^flox/flox^; *Trp53*^flox/flox^; *Rbl2*^flox/flox^ (*RPR2*) form the basis of a model of SCLC that recapitulates the salient genetic and histopathological features of this disease [22]. In *RPR2* mice, adenoviral vector transduction of lung epithelial cells with Cre recombinase deletes *Rb1*, *Trp53*, and *Rbl2,* which results in multifocal aggressive SCLC after ∼28 weeks [22, 23]. We replaced the *p53^flox^* allele in the *RPR2* model with the *p53*^XTR^ allele to generate an *Rb1^flox/flox^*; *Trp53^XTR/XTR^*; *Rbl2^flox/flox^* SCLC mouse model (*RP^XTR^R2*). *Trp53*^XTR^ allows for Cre-dependent inversion of the integrated gene trap to inactivate *p53* gene expression (*p53^TR^*)— a state that is marked by expression of EGFP, which is integrated into the gene trap. Once tumors are established, *p53* gene expression can be restored via a FlpO-dependent deletion of the gene trap (*p53^R^*) (Figure 1A). To control FlpO recombinase activity, we incorporated a *Rosa26^FlpO-ER^* allele into the *RP^XTR^R2* model. We endotracheally delivered adenoviral CMV-Cre (Ad:CMV-Cre) to a cohort of *RPR2* and *RP^XTR^R2* mice to induce the simultaneous inactivation of endogenous *Rb1*, *Trp53*, and *Rbl2* (Figure 1B). The resultant *RPR2* and *RP^TR^R2* tumors were indistinguishable in size, overall burden, and histological appearance (Figures 1C-1E). All tumors had small round cells with hyperchromatic nuclei, high nuclear:cytoplasm ratios, and expressed neuroendocrine markers (*e.g.* ASCL1, CGRP, UCHL1), consistent with established features of SCLC (Figure 1C) [24].

**Fig. 1.**
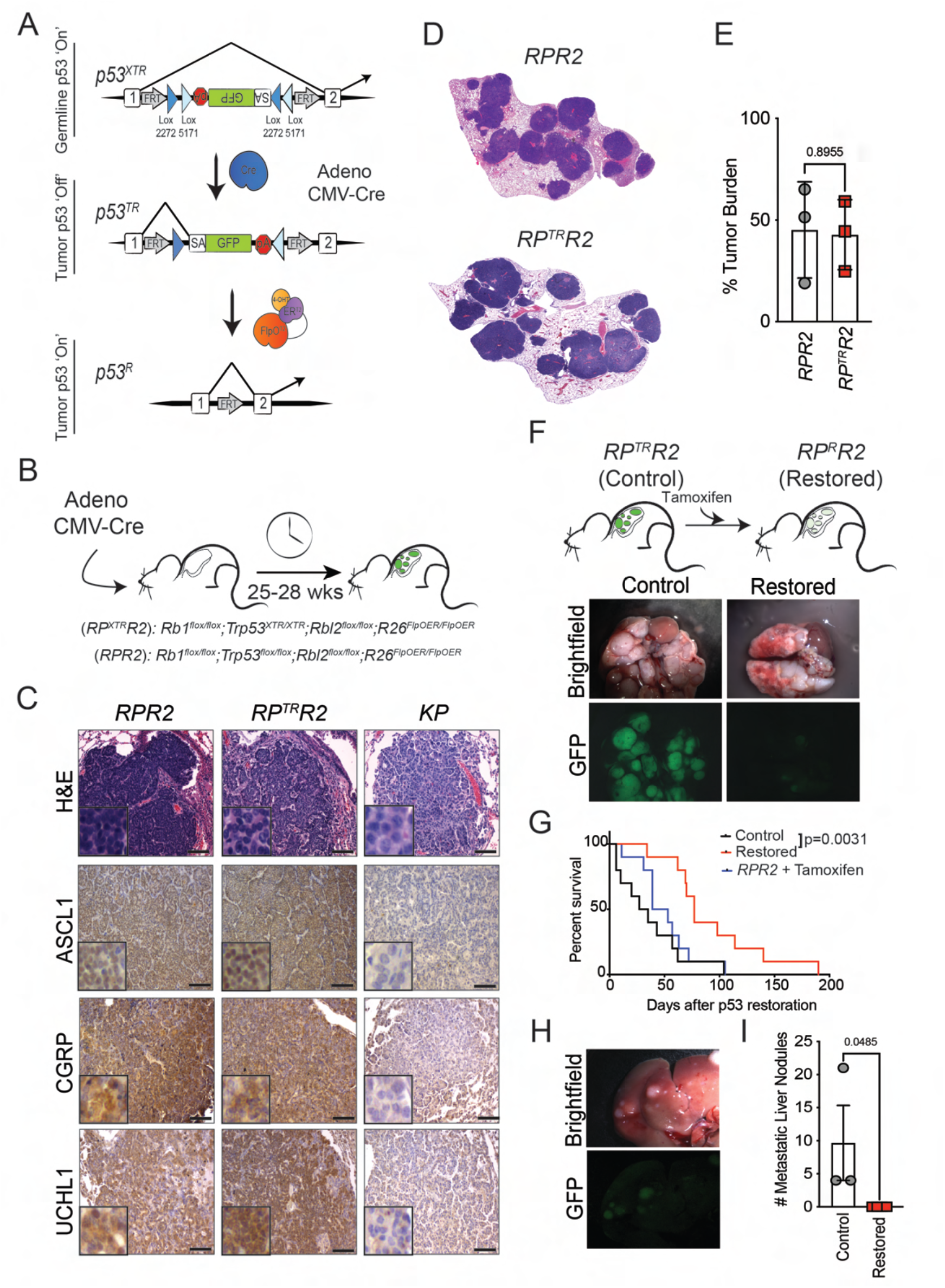
p53 restoration limits SCLC growth and metastasis *in vivo.* (**A**) Schematic of the *p53^XTR^* allele. Top, *p53^XTR^* (expressed): XTR gene trap cassette consists of a splice acceptor (SA), enhanced Green Fluorescent Protein (GFP), and the polyadenylation transcriptional terminator (pA). Stable inversion is achieved by the use of two pairs of mutually incompatible mutant *loxP* sites (Lox2272 and Lox5171) arranged in the ‘double-floxed configuration. Middle, *p53^TR^* (trapped): inhalation of Cre-expressing adenoviral vectors induces permanent gene trap inversion and conversion to the *p53^TR^* allele inactivating *Trp53* expression. Transcripts are spliced from exon 1 to the GFP reporter. Bottom, *p53^R^* (restored): the *Rosa26^FlpO-ERT2^* allele enables tamoxifen-dependent conversion of trapped *p53^TR^* to its restored *p53^R^* allelic state via excision of the gene trap. (**B**) *RPR2* and *RP^XTR^R2* mice were infected with 1.0×10^8^ p.f.u of Ad:CMV-Cre to initiate tumor formation. Between 25-28 weeks after tumor initiation, mice were treated with vehicle or tamoxifen, as described. (**C**) H&E and IHC for the neuroendocrine markers, ASCL1, CGRP, and UCHL1 in *RPR2*, *RP^TR^R2* and *KP* mice. Scale bars: 25µm for H&E, 25µm for IHC; insets are magnified 5×. (**D**) Representative scans of tumor-bearing lobes from *RPR2* or *RP^TR^R2* mice. (**E**) Tumor burden 25-28 weeks after tumor initiation. Statistical significance was determined by Student’s *t*-test. Error bars represent mean ± s.d. (**F**) Brightfield and fluorescent micrographs of lungs from *RP^TR^R2* mice treated with vehicle (Control) or tamoxifen (Restored). (**G**) Kaplan-Meier survival analysis. *RP^TR^R2*, n=10 mice per treatment group (Control or Restored); *RPR2*, n=10 mice. Statistical significance determined by Mantel-Cox log rank test. (**H**) Brightfield and fluorescent micrographs of metastatic liver nodules from control *RP^TR^R2* mice. (**I**) Quantification of metastatic liver nodules in control (n=3 mice) and restored (n=3 mice) after 9-16 days of tamoxifen treatment. Statistical significance determined by Student’s *t-*test.

To determine the impact of restoring p53 expression in SCLC, we administered tamoxifen starting 25-28 weeks after tumor initiation (Figure 1F). Consistent with the design of the *Trp53*^XTR^ allele, *RP^TR^R2* tumors expressed EGFP, whereas tumors from tamoxifen-treated animals (*RP^R^R2*) lost EGFP expression. Reactivation of p53 expression significantly enhanced overall survival in *RP^R^R2* animals compared to vehicle control-treated *RP^TR^R2* cohorts or tamoxifen-treated *RPR2* cohorts (Figure 1G). Moreover, metastatic nodules in the liver, a common site of metastatic spread in human SCLC were significantly fewer in *RP^R^R2* animals (Figures 1H, 1I) [25]. Notably, *RP^R^R2* animals eventually succumbed to SCLC burden, but these tumors were GFP+ indicating that p53 restoration failed in a subset of cells which then expanded (Figure S1). Nevertheless, the extended survival after p53 restoration suggested that p53 limits SCLC growth or disease progression.

### p53 restoration induces either senescence or necrosis in SCLC *in vivo*

To gain further insight into how p53 restoration influences tumor maintenance, we tracked tumor size over time using micro computed tomography (μCT). We observed that the mice with restored p53 expression had significantly less total volume of lung tumors relative to control animals after two weeks of restoration (Figure 2A). Individual tumors in control mice all increased in volume over the two-week observation period (1.3% to 378% growth). However, p53 restoration induced a heterogenous response with approximately half have limited growth relative to controls (3.6% to 98% growth) while the other half regressed (7.2% to 59.5% reduction) (Figure 2B). These data suggest that SCLC may have significant inter-tumor heterogeneity that impacts the potency of tumor suppression and/or the selection of specific tumor suppressive programs induced by p53 restoration.

**Fig. 2.**
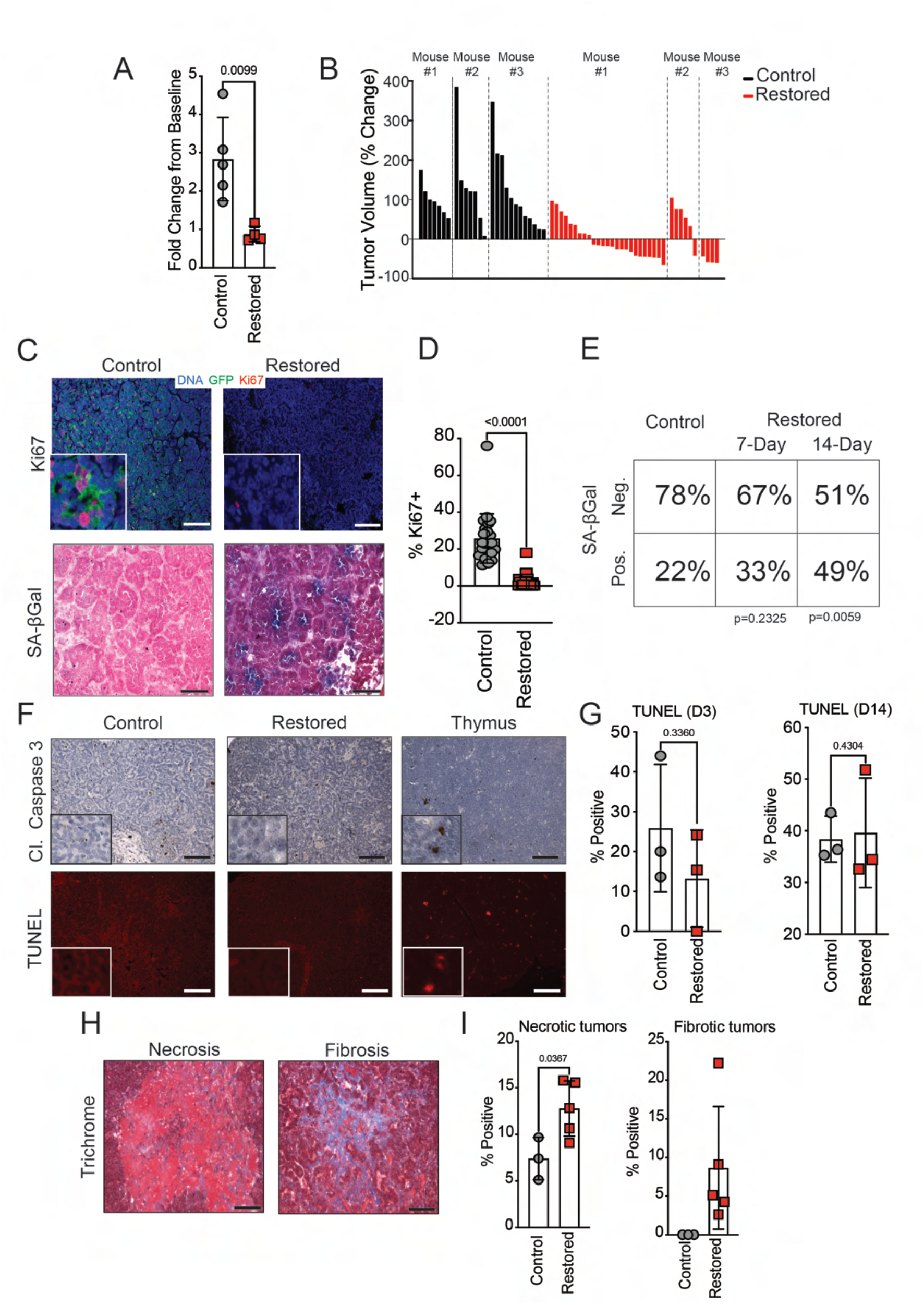
p53 restoration induces either senescence or necrosis in SCLC *in vivo.* (**A**) Tumor burden analysis from µCT scans. Fold change from baseline measurements. Control includes (*RPR2* (n=3) and *RP^TR^R2* (n=2)). Restored *RP^R^R2* (n=5) mice after 11-14 days of treatment. Statistical significance determined by Student’s *t*-test. Error bars represent mean ± s.d. (**B**) Waterfall plot of individual tumor volume plotted as percentage change from baseline in control (n=3 *RP^TR^R2*) and restored (n=3 *RP^R^R2*) mice. (**C**) Photomicrographs for GFP and Ki67 immunofluorescence and SA-β-Gal stained *RP^TR^R2* tumors 7 days after vehicle (Control) or tamoxifen (Restored) treatment. Scale bars: 25µmin for SA-β-Gal, 50µm for IF; insets are magnified 5×. (**D**) Quantification of Ki67 staining from (**c**). Statistical significance determined by Student’s *t-*test. Error bars represent mean ± s.d. (**E**) Contingency analysis of SA-β-Gal staining from (**c**); n=63 tumors from 7 Control mice; n=63 tumors from 4 Restored mice at t=7 days; n= 41 tumors from 4 Restored at t=14 days. Statistical significance determined by Fisher’s exact test. (**F**) Photomicrographs for cleaved Caspase-3 IHC or TUNEL stained *RP^TR^R2* tumors. Scale bars: 25µm for IHC, 50µm for TUNEL; insets are magnified 5×. (**G**) Quantification of TUNEL staining from (**f**) 3 or 14 days after tamoxifen treatment; n=3 mice for all treatment groups. Statistical significance was determined by Student’s *t*-test. Error bars represent mean ± s.d. (**H**) Representative photomicrographs for necrosis or fibrosis positive *RP^TR^R2* tumors from trichrome stained tissue sections. Scale bars, 25µm. (**I**) Quantification of trichrome staining from (**h**) after 14 days of tamoxifen treatment. n=3 mice for control mice; n=5 mice for restored mice. Statistical significance determined by Student’s *t*-test. Error bars represent mean ± s.d.

Most notable amongst p53-controlled tumor suppressive programs are senescence and apoptosis. Both of these fates have previously been observed in cancer models with reversible p53 inactivation [15–19]. Consistently, we found that p53 reactivation induces a significant decrease in cell proliferation across all tumors but leads to the progressive accumulation of senescence-associated-β-galactosidase (SA-β-Gal) activity in only a subset (Figures 2C-2E). However, p53 reactivation did not increase markers of apoptosis after 3 or 14 days, despite significant tumor regression (Figures 2F and 2G). Instead, we found a preponderance of necrotic and fibrotic areas specifically in a subset of SCLC tumors two weeks after p53 reactivation (Figures 2H, 2I). These findings suggest the possibility of inter-tumor heterogeneity wherein p53 reactivation in some tumors leads to the induction of cellular senescence and in others a necrotic form of cell death.

Senescence following p53 reactivation is associated with an inflammatory process that recruits immune cells ostensibly to eliminate cancer cells in hepatocellular carcinoma and Kras-driven lung adenocarcinoma [17, 20]. To establish the extent to which immune infiltration promotes tumor regression, we profiled immune cells by flow cytometry in SCLC tumors after p53 restoration. Although widespread immune infiltration into SCLC tumors after p53 reactivation was not prominent, there was a significant increase in F4/80+ macrophages after p53 restoration (Figure S2A). However, these macrophages were not localized within senescent tumors *per se* but were predominantly localized within tumors that had large necrotic and/or fibrotic regions after p53 reactivation. This suggests that these cells are likely infiltrating SCLC tumors to repair damaged tissue and clear dead cells, instead of directly culling senescent tumor cells (Figure S2B).

### SCLC-derived cell lines recapitulate heterogenous responses to p53 reactivation

To gain mechanistic insight into the distinct tumor suppressive programs induced by p53 in SCLC, we generated cancer cell lines from individual tumors from *RP^TR^R2* mice prior to p53 restoration (Figure 3A). Consistent with their SCLC origin, all cell lines expressed the neuroendocrine marker ASCL1 and lacked expression of RB (Figures S3A and S3B). Exposure of these cells to 4-hydroxytamoxifen (4-OHT) to activate FlpO-ER activity led to the conversion of the *Trp53^TR^* allele to the *Trp53^R^* state (Figure S3C). Likewise, EGFP reporter gene expression was diminished in concordance with the emergence of p53 protein expression and expression of the canonical p53-target gene p21 (Figure S3D and Figure 3B). Most importantly, and consistent with our *in vivo* findings, p53 reactivation induced two disparate effects across distinct cell lines (Figure 3C). A subset (4 of 8, 50%) of the cell lines remained viable after p53 reactivation (hereafter “Type V”) but halted the cell cycle and induced SA-β-Gal activity (Figures 3F-H, Figures S3E and S3F). In contrast, the remaining subset of cell lines (4 of 8, 50%) underwent a sudden cell lysis (necrosis) approximately 3 to 5 days after p53 reactivation (hereafter “Type D”) (Figures 3C-3E, Supplementary Video 1). Similar to our *in vivo* observations, we detected no markers of apoptosis in any of the Type D or Type V cell lines (Figure 3I, 3J). Importantly, cell death was not induced by potential off-target effects of tamoxifen treatment, or differences in p53 expression and/or stabilization (Figures S3G and S3H). Interestingly, in an allograft model, p53 reactivation in Type D cells induced varying degrees of tumor regression and necrosis, similar to what we observed in a subset of the autochthonous SCLC tumors (Figures 3K and 3L). These observations support the notion that SCLC tumors have tumor-intrinsic heterogeneity wherein different subclasses of SCLC tumors p53 induces disparate cancer cell-autonomous processes: senescence in Type V tumors and necrotic cell death in Type D tumors.

**Fig. 3.**
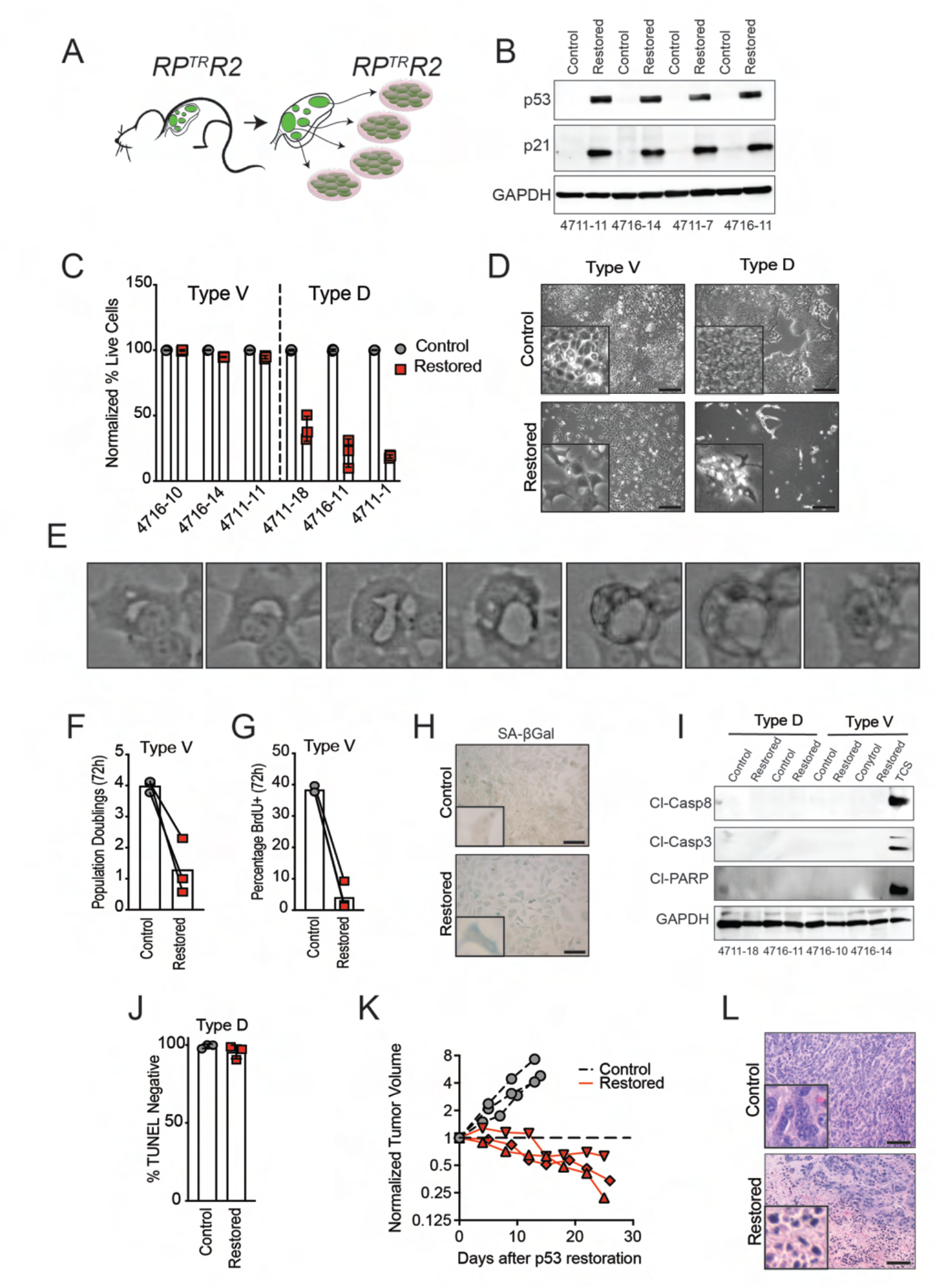
p53 reactivation induces cellular senescence or non-apoptotic cell death in a cancer cell autonomous manner. (**A**) *RP^TR^R2* mice were infected with 1.0×10^8^ p.f.u of Ad:CMV-Cre to initiate tumor formation. Between 25-28 weeks after tumor initiation, tumors were excised for cancer cell line establishment. (**B**) Immunoblot analysis of representative *RP^TR^R2* cell lines (n=4) for p53 and p21 expression 24hrs after 4-OHT treatment. GAPDH is a loading control. (**C**) Flow cytometry-assisted cell viability assay in representative Type V (n=3) and Type D (n=3) cells 5 days after 4-OHT treatment. Live cell percentage determined by quantification of DAPI negative population; n=3 technical replicates for all cell lines and treatment groups. Error bars represent mean ± s.d. (**D**) Brightfield photomicrographs from Type V and Type D cell lines after 5 days of 4-OHT treatment. Scale bars, 250µm. (**E**) Time lapse video of a Type D cell dying after p53 reactivation. Video accounts for 72-hour time span between 24 and 96 hours after p53 restoration.(**F**) Quantification of cell population doublings 72 hours after 4-OHT treatment. Each symbol represents the mean of n=3 technical replicates per cell line. (**G**) Quantification of BrdU+ cells 72 hours after 4-OHT treatment. Symbols represent independent Type V cell lines (n=3). (**H**) Brightfield photomicrographs of SA-β-Gal stained Type V cells representative of three independent Type V cell lines. Scale bars, 25µm; insets are magnified 5×. (**I**) Immunoblot analysis for apoptosis markers in Type V and Type D cells 4 days after 4-OHT treatment. Positive control (TCS) treated with TNF-α (1µg/mL), Smac mimetic (SM-164; 100nM), and cyclohexamide (10µg/mL) for 8 hours. GAPDH is loading control. (**J**) Quantification of flow cytometry-assisted TUNEL staining in Type D cells. Symbols represent independent Type D cell lines (n=3). Error bars represent mean ± s.d. (**K**) Representative growth curve from Type D allograft tumor(s). Each line represents replicate (n=3) allograft of one Type D cell line. Experiment was conducted in at least n=3 Type D cell lines with n=3 replicates per treatment group. (**L**) Brightfield photomicrographs of allograft tumors generated from Type D cell line used in (**j**) 7 days after tamoxifen treatment. Each condition had n=2 tumors. Scale bars, 25µm.

### p53 induces a novel form of cyclophilin-dependent necrosis in SCLC

p53 activity has been linked to induction of distinct, regulated forms of cell death [26–30]. To gain insight into the mechanism of cell death induced by p53 reactivation in SCLC, we developed mCherry-expressing and CRISPR-based vectors (LentiCRISPRv2-mCherry) targeting critical components of distinct cell death pathways (Figure 4A). These vectors allow us to conduct positive enrichment assays, where knockout of critical death effector molecules would lead to enrichment of mCherry-positive cells after p53 reactivation relative to controls. CRISPR-mediated knockout of key apoptosis (*Bax, Bbc3),* necroptosis (*Mlkl, Ripk1*) or autophagy *(Atg5)* regulators did not impact p53-mediated death (Figure 4B). We then treated Type D cells with pharmacological inhibitors of apoptosis (ZVAD-FMK), ferroptosis (Ferrostatin-1), necropstosis (Necrostatin-1s), and mitochondrial permeability transition pore (mPTP)-induced necrosis (Cyclosporin A). Surprisingly, Cyclosporin A (CsA) treatment protected all Type D cell lines from p53-mediated cell death (Figure 4C and S4A,B). mPTP-induced necrosis has been attributed to a non-transcriptional function of p53 where it binds cyclophilin D (encoded by *Ppif*) to disrupt mitochondria membrane permeability and induce necrosis during tissue ischemia [26]. However, we could not establish p53 localization to the mitochondria or detect evidence for a unique impact on mitochondria physiology after p53 restoration in Type D SCLC cell lines (Figures S5). Moreover, CRISPR-mediated inactivation of *Ppif* did not block p53-mediated cell death in Type D SCLC. These data suggest that although CsA potently blocks p53-mediated cell death in Type D SCLC, the mechanism is distinct from mPTP-induced necrosis.

**Fig. 4.**
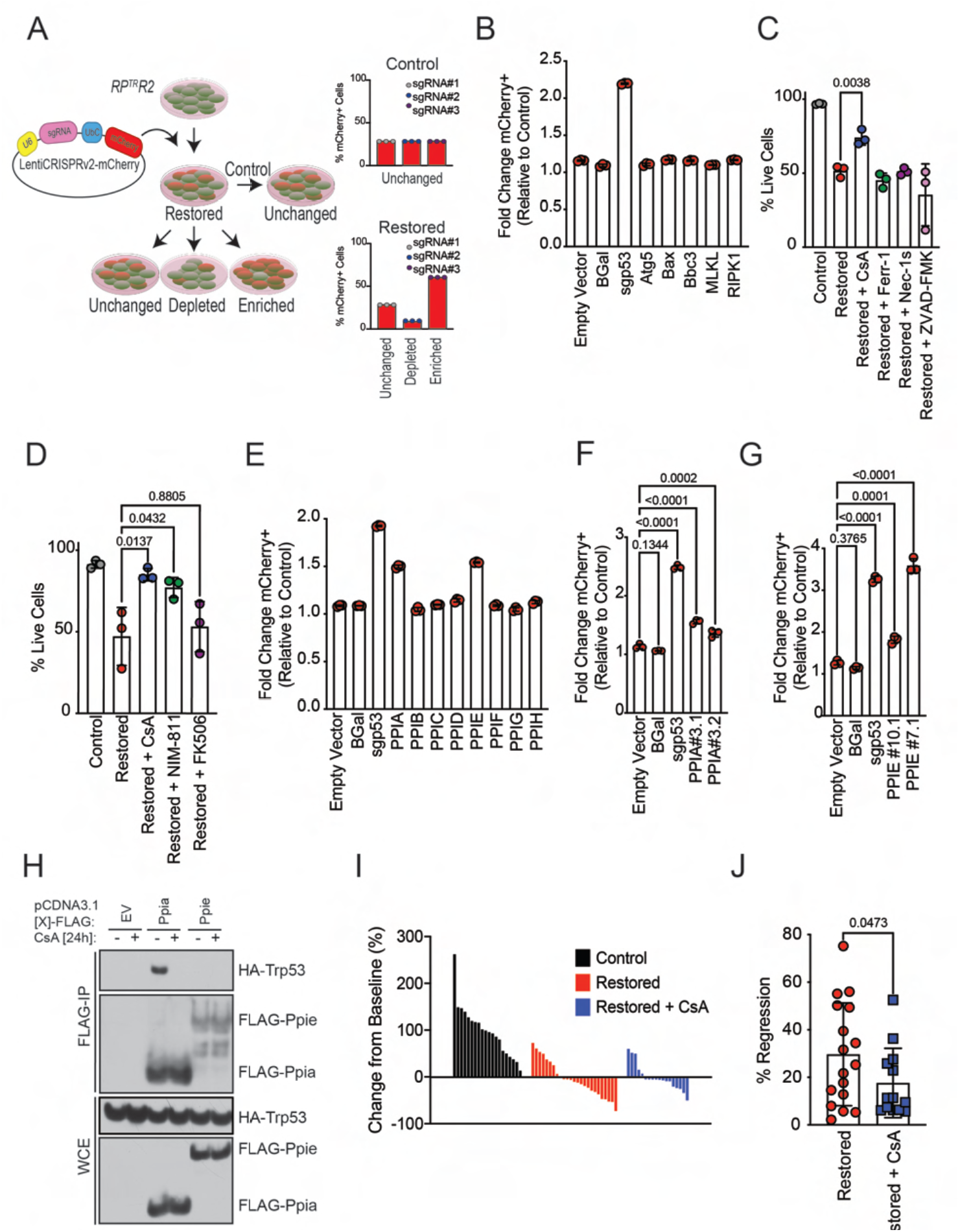
p53-mediated Type D cell death is Cyclophilin-dependent. (**A**) Schematic of LentiCRISPRv2-mCherry transduction of Type D cells for positive selection assay. Type V cells are infected with mCherry-expressing lentiviral vectors expressing Cas9 and sgRNAs targeting specific cell death mediators or cyclophilins. Baseline mCherry expression is measured in viable (DAPI-) control cells using flow cytometry. Upon p53 restoration, Changes in the proportion of mCherry-positive cells quantified in live (DAPI-) cells. Data plotted as fold change mCherry+ population relative to Control (vehicle treated) cells. (**B**) Fold change in the proportion of mCherry-positive cells 72hrs after 4-OHT treatment in a representative Type D cell line expressing sgRNAs targeting distinct apoptosis (*Bax, Bbc3*) and necroptosis (*Mlkl, Ripk1*) regulators. Empty Vector, β-Gal, and a p53 targeting sgRNA used as controls. Each symbol represents a technical replicate (n=3). Error bars represent mean ± s.d. Conducted in n=2 independent Type D cell lines. (**C**) Type D cells were treated with 4-OHT and specific cell death inhibitors for 72hrs. Percentage of live cells (DAPI-) were determined using flow cytometry. Each symbol represents a technical replicate (n=3). Statistical significance was determined by Student’s *t-*test. Error bars represent mean ± s.d. Experiment was conducted in n=2 independent Type D cell lines. (**D**) Representative Type D cell line was treated with 4-OHT and cyclophilin (CsA, NIM-811) or FKBP (FK506) inhibitors for 72hrs. Percentage of live cells (DAPI-) were determined using flow cytometry. Each symbol represents mean of technical replicates (n=3). Statistical significance was determined by Dunnett’s multiple comparison test. Error bars represent mean ± s.d. Data represent n=3 independent experiments. (**E**) Fold change in the proportion of mCherry-positive cells 72hrs after 4-OHT treatment in representative Type D cell line expressing sgRNAs targeting distinct cyclophilins. Empty Vector, β-Gal, and a p53 targeting sgRNA used as controls. Each symbol represents a technical replicate (n=3). Error bars represent mean ± s.d. Experiment was conducted in n=2 independent Type D cell lines (Figures 4E and 4F, S3E). (**F,G**) Validation of results in E using an independent Type D cell line and additional sgRNAs targeting exons 3 of the *Ppia* gene (F) or exons 7 and 10 of the *Ppie* gene (G). Each symbol represents a technical replicate (n=3). Statistical significance was determined by Dunnett’s multiple. comparison test. Error bars represent mean ± s.d. (**H**) Immunoblot for PPIA, PPIE and p53 expression 24hrs after CsA treatment. Whole cell extracts (WCE) or immunoprecipitated (FLAG-IP) samples from HEK293T cells overexpressing FLAG-*Ppia* or FLAG-*Ppie,* and HA-*Trp53.* (**I**) Individual tumor volume analysis using µCT. Waterfall plot percentage change from baseline in control (n=3, n=20 tumors), restored (n=3, n=25 tumors), and restored + CsA (n=5, n=18 tumors) mice after 15 days of treatment. (**J**) Quantification of percentage regression of tumors in (I). n=17 regressing tumors from n=3 mice treated with tamoxifen; n=13 regressing tumors from n=3 mice treated with tamoxifen and CsA. Error bars represent mean ± s.d. Statistical significance determined by Student’s *t*-test.

CsA is most notable for its ability to suppress T cell function by forcing a neo-protein-protein interaction with calcineurin and cyclophilin, which blocks downstream NFAT signaling [31–33]. FK506 is another small molecule that forces a distinct neo-protein-protein interaction of FKBP12 with calcineurin which also blocks NFAT signaling [32, 34, 35]. To distinguish whether this p53-mediated death requires NFAT activation or an alternative activity of cyclophilins themselves, we treated cells with a cyclophilin-specific inhibitor that does not affect calcineurin/NFAT signaling (NIM-811) or FK506 which blocks calcineurin/NFAT but not cyclophilin activity [32, 34, 35]. NIM-811, but not FK506, blocked cell death similarly to CsA, confirming that cyclophilins are key effectors of p53-induced death in Type D cells (Figure 4D, Figures S4C).

To establish the role of specific cyclophilin(s) in p53-mediated necrosis in Type D cells, we generated LentiCRISPRv2-mCherry vectors that targeted all cyclophilin family members known to interact with CsA (*Ppia, Ppib, Ppic, Ppid, Ppie, Ppif, Ppig, Ppih*)[36]. Using the same mCherry/CRISPR enrichment assay described above (Figure 4A), we discovered that *Ppia* and *Ppie* sgRNAs were significantly enriched upon p53 reactivation (Figures 4E-4G). On the other hand, sgRNAs targeting other cyclophilin family members, including *Ppif*, did not block p53-mediated death (Figures 4E and S4D). This finding suggests that Cyclophilin A and Cyclophilin E are critical determinants of p53-mediated death in Type D cells. Cyclophilin A (encoded by *Ppia*) has been shown to physically interact with p53 to affect p53 target gene selection [37]. We performed co-immunoprecipitation experiments using FLAG-PPIA or FLAG-PPIE, and HA-p53. Consistently, Cyclophilin A strongly interacted with p53 (Figure S4E). However, we observed no interaction between Cyclophilin E (encoded by *Ppie*) and p53 (Figure S4E). Interestingly, deletion of the proline rich domain of p53 decreased binding of Cyclophilin A to p53 and the additional deletion of the DNA binding domain of p53 completely abrogated the interaction (Figure S4F). To elucidate whether CsA modulates p53-cyclophilin interactions, we co-immunoprecipitated FLAG-PPIA or FLAG-PPIE, and HA-p53 in the presence or absence of CsA. Strikingly, the CypA-p53 interaction was completely inhibited by CsA treatment (Figure 4H). These data suggest that Cyclophilin A could be directly modulating p53 function and that disrupting the CypA-p53 interaction with CsA could compromise the ability of p53 to induce cell death in Type D SCLC.

To establish the role of cyclophilins in p53-mediated death *in vivo*, we tracked individual tumor size in live mice over time using μCT after daily CsA (15mg/kg) treatment. Consistent with our *in vitro* findings, cyclophilin inhibition diminished tumor regression *in vivo,* suggesting that CsA treatment suppresses p53-mediated death (Figures 4I and 4J). Taken together, these data reveal that p53 induces a novel form of cell death dependent on Cyclophilin A and Cyclophilin E in SCLC, and that inhibition of cyclophilins can limit p53-mediated tumor cell death *in vitro* and *in vivo*.

### Type V and Type D cells have features of distinct molecular subtypes of human SCLC

Cellular context influences p53 function to induce distinct tumor suppressive programs [38–42]. Thus, we conducted RNA-sequencing in Type D and Type V SCLC cells to gain insight into context-specific differences that may differentially regulate p53 function. Principal component analysis (PCA) revealed that Type V or Type D class is the primary distinguishing feature among all samples, making up >47% of the variation in the data (Fig. 5A). Secondly, p53 restoration status made up >12 % of the variation in the data (Control versus Restored). To further characterize the baseline transcriptional differences (Control vs Control) between Type D and Type V SCLC cells, we conducted differential gene expression and gene set enrichment analysis (GSEA) [43, 44]. While only 109 genes were enriched in Type D cells and 57 genes were enriched in Type V cells (Figure 5B), multiple gene sets distinguished Type V and Type D cells. Notably, Type D cells were enriched for metabolic gene signatures, whereas Type V cells were enriched for gene signatures regulating epithelial mesenchymal transition, Wnt and hedgehog signaling, and inflammatory processes (Figure 5C). Human SCLC is a highly heterogeneous disease that has recently been stratified into four molecular subtypes based on expression of lineage defining factors: ASCL1 (SCLC-A), NEUROD1 (SCLC-N), YAP1 (SCLC-Y), POU2F3 (SCLC-P) [4]. As such, we determined whether Type D and Type V cells were enriched for human SCLC-subtype-specific gene signatures [45].

**Fig. 5.**
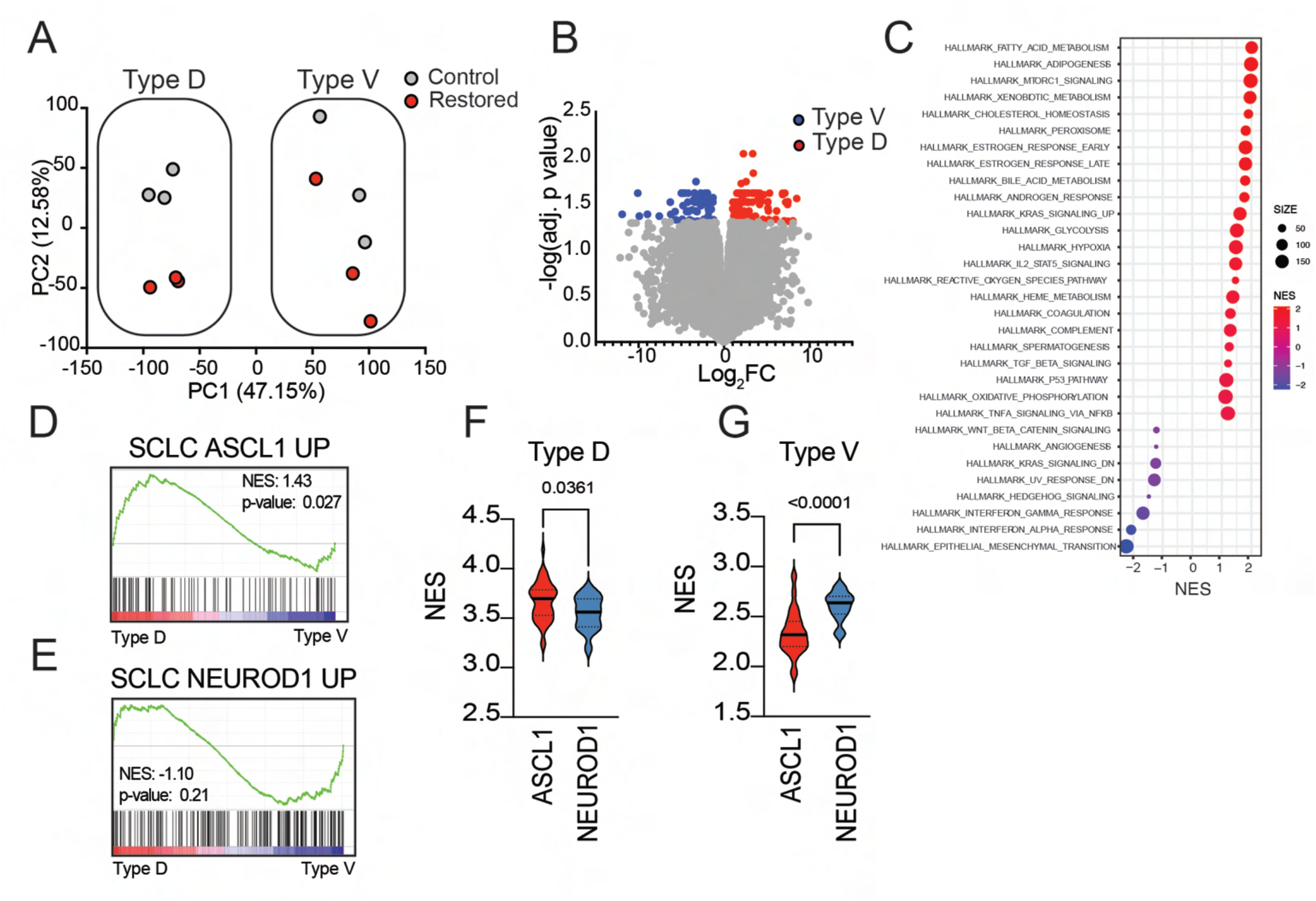
Type V and Type D cells have features of distinct SCLC subtypes. (**A**) Principal component analysis of Type D (n=3) and Type V (n=3) cell lines 72 hours after treatment with vehicle (Control) or 4-OHT (Restored). (**B**) Volcano plots of RNA-sequencing data indicating in untreated Type D and Type V cells. Colored dots represent genes that are differentially enriched (log_2_ fold-change greater-than 1 and false discovery rate (FDR)-adjusted *P*-value less-than 0.05) in Type D (n=109, red) or Type V (n=57, blue) cells. **(C)** Bubble plot representation of all GSEA hallmark gene sets with an FDR-adjusted *P-*value lower than 0.25 in untreated Type D (red) or Type V cells (blue). Normalized enrichment score (NES) plotted and the “SIZE” of each dot represents the number of genes within the gene set. (**D,E**) GSEA of “SCLC_ASCL1_UP” (**d**) and “SCLC_NEUROD1_UP” (**e**) gene signatures from (Ireland et al., 2020) in untreated Type D and Type V cells. (**F,G**) Violin plots indicating normalized enrichment scores of ‘Type D’ (F) and ‘Type V’ (G) gene signatures in human SCLC-ASCL1 (n=47) and SCLC-NEUROD1 (n=15) cell lines. Single sample GSEA analysis was performed to quantify enrichment of specific gene signatures in gene expression datasets from human SCLC cell lines obtained using CellMiner-SCLC (Tlemsani et al., 2020). Statistical significance was determined by Student’s *t*-test.

Interestingly, Type D cells were significantly enriched for the “classic” SCLC-A molecular subtype, whereas Type V cells were moderately enriched for the “variant” SCLC-N molecular subtype (Figures 5D and 5E). To further validate this finding, we generated gene signatures associated with Type D and Type V cells based on differentially regulated genes (Figure 5B) and used SCLC-CellMiner to compare our signatures to human SCLC cell line gene expression datasets [46]. Consistent with our results, SCLC-A human cells are significantly enriched for the Type D signature, whereas SCLC-N human cells are enriched for the Type V signature (Figures 5F and 5G). Taken together, Type D and Type V SCLC are defined by basal gene expression related to two molecular subtypes of human SCLC.

### SCLC heterogeneity does not dictate p53 binding

To examine p53 DNA binding patterns, we conducted chromatin immunoprecipitation (ChIP)-sequencing 48 hours after p53 reactivation in Type D and Type V cells. As a proof of principle, we observed high p53 ChIP-signal (Restored) and very low noise (Control) at the prototypical p53 target gene, *Cdkn1a* (Figure S6A). Across the genome, the location of p53 binding was largely similar across all SCLC cell lines; however, Type V cells had a significantly higher degree of p53 binding at most sites (Figures S6B-S6D). Differential peak calling analysis identified 1,032 peaks in Type V cells which corresponded to 508 differentially bound genes (Type V DBGs). Only 3 peaks had significantly higher binding in Type D cells, each of which corresponded to a specific gene (Type D DBGs) (Figure S6E). However, differential peak calling between Type D and Type V cells did not reflect distinct changes in gene expression after p53 restoration. In fact, 434 out 508 Type V DBGs were not differentially expressed at all after p53 restoration in Type V or Type D cells. Moreover, for those 74 Type V DBGs that were differentially expressed after p53 restoration, the vast majority were also similarly up- or down-regulated in Type D cells (Figures S6F-H). Expression of the 3 Type D DBGs was likewise not distinctly up- or down-regulated by p53 in Type D or Type V cells (Figures S6I). These data suggest that differential p53 binding is not a robust predictor of the distinct physiological programs orchestrated by p53 in Type D and Type V SCLC and that p53 binds to promoter-proximal sequences within a similar repertoire of genes in both Type D and Type V cells.

### p53 induces a distinct transcriptional program in Type D SCLC that is cyclophilin-dependent

We conducted differential gene expression analysis between control and p53-restored Type D and Type V cells to determine whether p53 regulated distinct transcriptional programs in each. We identified 764 genes that were differentially expressed (p<0.05 and log_2_ fold change ≥ 1) after p53 reactivation in Type D cells and 250 genes that were differentially expressed in Type V cells 72 hours after p53 restoration (Figures 6A and 6B). Importantly, *Trp53* transcript abundance and the induction of p53 Hallmark Genes were similarly up-regulated after p53 restoration, suggesting that regulation of the *Trp53* gene *per se* or p53’s regulation of canonical target genes were not responsible for disparate cell fates after p53 restoration in Type D and Type V cells (Figures 6C and 6D). Taking the union of the p53-regulated genes identified in Type D and Type V cells, we performed unbiased hierarchical-clustering analysis in control and p53-restored samples. The genes clustered into 6 distinct gene expression modules (Supplementary Table 2). Generally, the expression patterns of all genes in each Module moved in the same direction after p53 restoration in both cell types regardless of whether the gene was identified in Type V or Type D samples (Figure 6E). However, the 102 genes in Module 5 were disproportionally elevated in Type D cells after p53 restoration compared to a negligible induction in Type V cells (Figures 6E). Interestingly, the 102 genes fell into multiple gene ontology (GO) and functional enrichment categories associated with regulation of cell death processes (Figure S7 and Supplementary Table 2). These data suggest that p53 may be distinctly regulating a small subset of genes to induce distinct tumor suppressive programs in Type D SCLC.

**Fig. 6.**
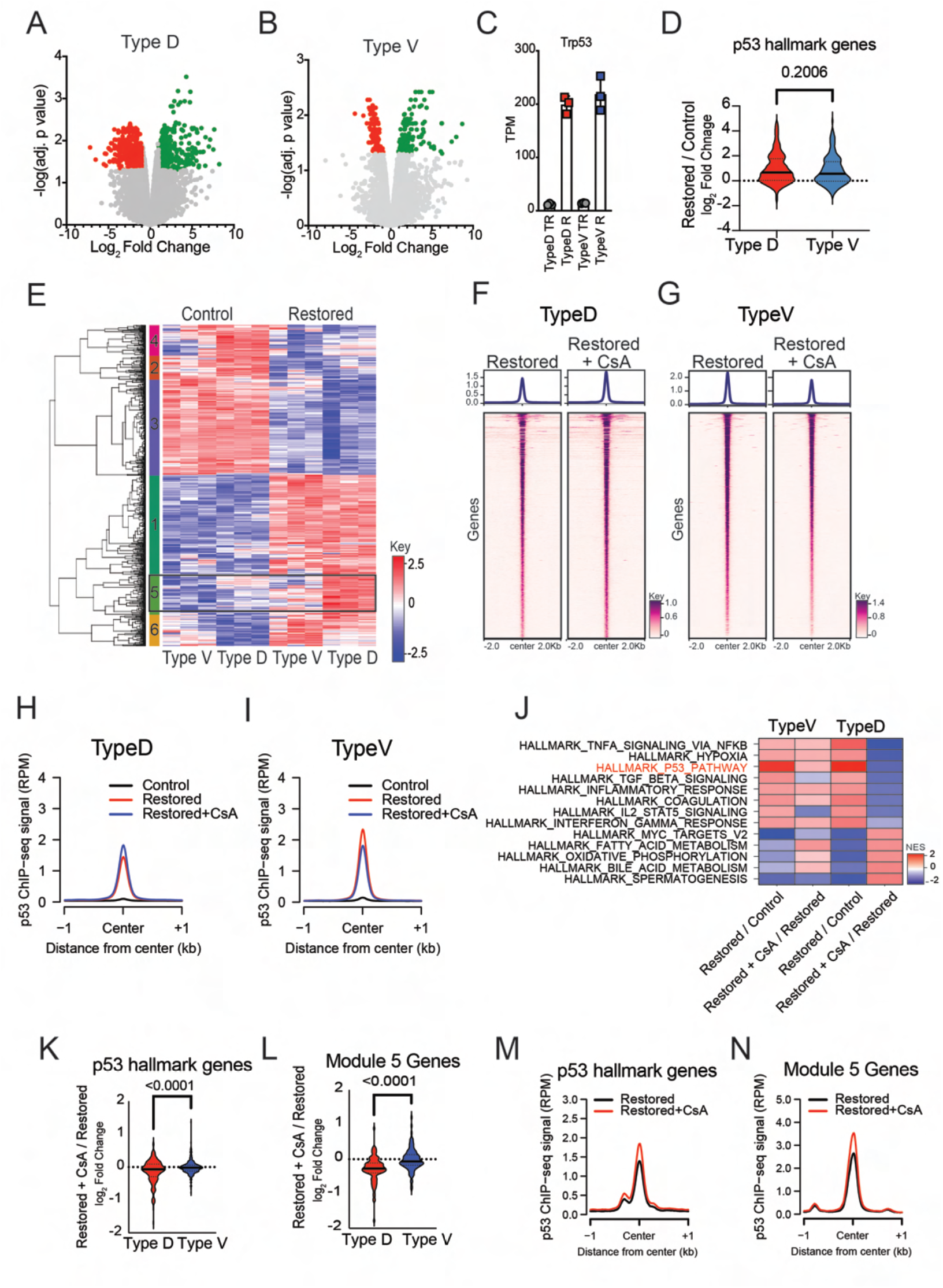
p53 controls a distinct transcriptional program in Type D SCLC that is dependent on cyclophilin activity. (**A,B)** Volcano plots of RNA-sequencing data indicating in Type D and Type V cells. Colored dots represent genes that are differentially enriched (log_2_ fold-change greater-than 1 and false discovery rate (FDR)-adjusted *P*-value less-than 0.05) in Restored (n=401, green) or Control (n=363, red) cells. (**C**) Quantification of transcripts per million (TPM) for *Trp53* in Control and Restored Type D (n=3) and Type V (n=3) cell lines. Error bars represent mean ± s.d.(**D**) Violin plots indicating log_2_ fold-change in expression of canonical p53 targets from the GSEA gene set ‘HALLMARK_P53_PATHWAY’ in Restored Type D (n=3) and Type V (n=3) cells. Statistical significance was determined by Student’s *t*-test. (**E**) Heatmap representation of the union of differentially expressed genes identified in (B,C) for Type D (n=3) and Type V (n=3) cells. Unbiased hierarchical clustering identifies 6 modules associated with distinct patterns of gene expression. The color key represents gene expression levels with darker colors representing higher (red) or lower (blue) gene expression. (**F,G**) Heatmap representation of Ch-IP seq analysis for p53 bound peaks 48hrs after 4-OHT and CsA treatment in Type D (n=3) (F) and Type V (G) (n=3) cells. Heatmaps are centered on p53-bound peaks across a ±2kb window. (**H,I**) Metagene analysis of p53 ChIP-seq signal between Control, Restored andRestored + CsA in Type D (n=3) (H) and Type V (n=3) cells (I). Data are centered on p53-bound peaks across a ±1kb window. (**J**) Heatmap representation of GSEA hallmark gene sets with an FDR-adjusted *P-*value lower than 0.25 from the comparisons listed in the labels along the y-axis. The color key represents normalized enrichment score (NES). (**K,L**) Violin plots indicating log_2_ fold-change expression of canonical p53 targets from the GSEA gene set ‘HALLMARK_P53_PATHWAY’ (K) or Module 5 genes (I) in Restored and Restored + CsA Type D (n=3) and Type V (n=3) cells. Statistical significance determined by Student’s *t*-test. (**M,N**) Comparison of p53 ChIP-seq signal between Restored and Restored + CsA Type D (n=3) cells for p53 hallmark genes (M) and Module 5 genes (N). Data are centered on p53-bound peaks across a ± 1kb window.

That cyclophilin activity is required for p53-mediated Type D SCLC cell death and that p53 interacts with Cyclophilin A in a CsA dependent manner led us to hypothesize that cyclophilins may be required for p53-dependent transcription in this context. Thus, we conducted ChIP- and RNA-sequencing experiments after p53 restoration while treating SCLC cells with CsA. Surprisingly, CsA treatment did not significantly influence p53 binding across the genome in Type D or Type V cells (Figure 6F-I). Moreover, CsA treatment did not alter gene expression prior to p53 restoration in any cell line, including expression of cyclophilin family members (Figure S8A-C). To explore the effect of CsA on p53-induced gene expression, we conducted GSEA and compared how CsA modulated p53 transcriptional output between Type D and Type V cells. Strikingly, CsA treatment abolished the enrichment of Hallmark p53 Pathway genes in Type D cells after p53 restoration. However, the effect of CsA on Hallmark p53 Pathway genes in Type V cells had the reverse effect, slightly promoting expression of these genes after p53 restoration (Figure 6J). Consistently, after p53 restoration, the average change in expression for each gene in the Hallmark p53 Pathway was significantly diminished in Type D cells in the presence of CsA treatment. However, the addition of CsA to Type V cells had no effect (Figure 6K). Given that Module 5 genes were predominantly induced by p53 restoration specifically in Type D cells (Figure 6E, Figure S7E), we determined the extent to which CsA treatment influenced p53-mediated transcription of these genes. Strikingly, CsA abolished the induction of Module 5 genes in Type D cells (Figure 6L). Despite the failure of p53 to induce canonical target gene expression generally and most potently at Module 5 loci in the presence of CsA, p53 bound equally well, if not slightly better, to promoter proximal regions within these genes (Figures 6M and 6N, Figures S8D and S8E). These data indicate that p53 transcriptional output, and not target gene binding, requires the additional activity of cyclophilins specifically in Type D SCLC. Moreover, p53 can transactivate a unique subset of genes (Module 5) specifically in Type D SCLC that are likely required for p53-mediated cell death in SCLC.

## Discussion

Our study provides key insight into the persistent selective requirement for p53 inactivation and the potential therapeutic responses to p53 reactivation in SCLC. Previous studies using mouse models with reversible p53 expression in distinct cancer contexts predominantly report a uniform response to p53 reactivation, with tumors either undergoing a senescence response followed by immune-mediated tumor clearance or apoptotic cell death [15–20]. We demonstrate that p53 reactivation limits SCLC disease progression by inducing senescence without a major immune reaction or by cyclophilin-dependent necrotic cell death in both autochthonously-arising tumors and tumor-derived cell lines. That p53 induces disparate tumor suppressive programs in SCLC suggests that distinct cellular contexts exist within tumors to modulate p53 function. By examining the basal transcriptional programs present in senescing (Type V) and dying (Type D) tumor-derived cells, we determined that each SCLC subtype is transcriptionally distinct. Importantly, Type V and Type D cells have gene expression programs that align most with the *ASCL1* and *NEUROD1* molecular subtype of human SCLC respectively. These lineage-defining transcription factors have been shown to regulate distinct genetic programs in SCLC and disparately regulate the expression of distinct oncogenic genes to drive tumor heterogeneity [47, 48]. Tumor initiation in our model was driven by adenoviral CMV-Cre, which can potentiate formation of SCLC tumors from distinct cells of origin that influence tumor evolution and metastatic mechanisms [10]. Thus, our findings warrant further investigation into the impact of cell-of-origin and driver mutations on p53-mediated tumor suppression in SCLC. These insights could identify context-dependent regulators of p53 function.

Peptidyl-proline isomerases (PPIAses) are a protein superfamily composed of cyclophilins, parvulins, and FK506-binding proteins [49]. PPIases canonically regulate protein function by inducing conformational changes after catalyzing the isomerization of proline residues [50, 51]. However, further structural and functional analysis of cyclophilin family members have identified non-canonical cyclophilin functions such as RNA-binding, spliceosome regulation, and U-box ubiquitin ligase activity [36, 52–55]. Unsurprisingly, this breadth of functions have implicated cyclophilins in the regulation of a plethora of biological processes and made them attractive therapeutic targets in the context of disease [26, 56–58]. Previous studies establish that p53 can directly interact with Cyclophilin D (*Ppif)* and Cyclophilin A (*Ppia)* to potentiate mitochondrial permeability pore (mPTP) opening and necrosis, or regulate cell cycle and apoptosis function, respectively [26, 37]. Our data implicate cyclophilins as key regulators of p53-mediated transcription and subsequent cell death program specifically in Type D SCLC cells. p53-mediated cell death is non-apoptotic and dependent on Cyclophilin A and Cyclophilin E for its execution. While it remains unclear how these cyclophilin family members impact p53 transcriptional output, Cyclophilin A can directly interact with p53 and this interaction is abolished by CsA. We speculate that the PPIase activity of cyclophilins may directly alter the efficiency of p53 transactivation or the recruitment of specific factors that promote expression of key regulators needed to execute the cell death program. Interestingly, Cyclophilin E did not interact with p53 directly in our analysis, suggesting a possible indirect mechanism for Cyclophilin E to modulate p53-mediated death. Future studies are required to determine how these cyclophilins modulate p53 function to induce cell death.

Cellular context influences the ability of p53 to induce specific tumor suppressive programs [38–41]. Moreover, p53 restoration in distinct cancer types was recently described to induce cancer type-specific p53 binding and transcriptional outputs that were likely responsible for differential induction of apoptosis in mouse lymphoma models, or senescence in sarcoma and lung adenocarcinoma models [42]. We profiled p53 binding and transcriptional output in Type D and Type V SCLC cells and observed that while global p53 binding was similar, p53 distinctly regulated a subset of genes between SCLC subtypes. Interestingly, although Module 5 genes are associated with diverse biological processes (*e.g* vesicular trafficking, cell growth), they were enriched for some genes specifically involved with canonical forms of apoptosis (*e.g. Apaf1 and Perp)* which may suggest some mechanistic overlap. However, the precise mechanism of p53-mediated cell death does not neatly fit into any defined cell death programs. That p53-mediated Type D SCLC cell death is abolished by cyclosporin A, and that cyclosporin A similarly abolishes the induction of a distinct set of genes specifically induced in Type D SCLC, strongly suggests p53 orchestrates a deliberate program of cell death. However, the program is not associated with mitochondrial dysfunction, activation of caspases, or extensive fragmentation of DNA, and is not inhibited by small molecules targeting established determinants of major forms of programmed cell death. Elucidating the molecular underpinnings of p53-mediated Type D SCLC will be a major focus of future work that may identify novel therapeutic targets to activate this latent form of cell death in SCLC.

In summary, our study shows that p53 reactivation in SCLC identifies two tumor subtypes (*Type V, Type D*) that either undergo senescence or a novel cyclophilin-dependent cell death, and identifies cyclophilins as context-dependent regulators of p53-mediated transcription (Figure 7).

**Fig. 7.**
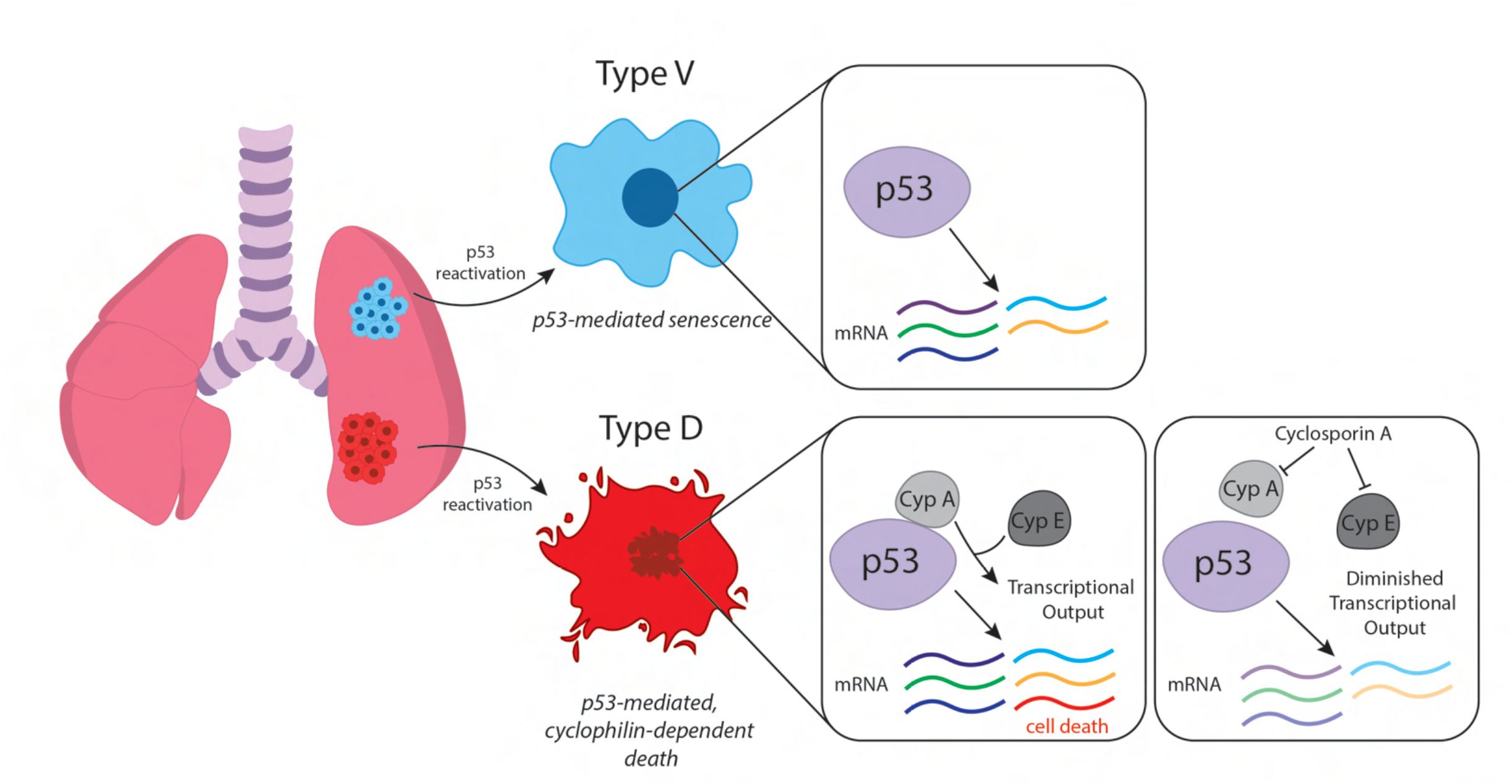
Model: p53 induces Cyclophilin-dependent cell death or senescence in distinct SCLC tumor subtypes. Reactivation of p53 in small cell lung cancer identifies two tumor subtypes that are suppressed by induction of distinct p53-mediated tumor suppressive programs. Type V tumors undergo senescence after p53 reactivation whereas, Type D tumors undergo a novel form of p53-mediated, cyclophilin-dependent cell death. p53 controls the expression of widely similar genes between tumor subtypes; however, p53 reactivation in Type D cells specifically induces the expression of select genes (Module 5) that are associated with cell death but whose induction is dependent on the activity of cyclophilin. Cyclophilins may potentiate p53 transcriptional output in a direct (Cyclophilin A) or indirect manner (Cyclophilin E). Cyclosporine A (CsA) blocks p53-mediated death likely by abrogating CypA-p53 interactions and inhibition of CypE function, each of which are required for efficient induction of p53-mediated cell death in Type D small cell lung cancer.

## Acknowledgements

We would like to thank ULAR staff for animal husbandry, the Molecular Pathology and Imaging Core (MPIC) for histological analysis and their technical expertise, E. Blankemeyer for computed tomography, D. MacPherson for *RP* cell line, I. Asangani and members of his lab for help with Next Generation Sequencing, and M. Murphy, M. Winslow, and members of the Feldser Laboratory for manuscript critique.

## Author Contributions

J.A., M.C., and K.M.A. performed animal studies. J.A. performed micro-CT analyses. J.A., N.M., H.M., M.C., and M.R.T., performed cell culture studies. A.C.G. performed Seahorse Mito Stress analysis. K.M.A and J.L performed immunological analysis with supervision from B.Z.S. J.A., Q.L., and N.F.F. performed bioinformatics analyses. L.W. supervised ChIP and bioinformatic analysis. G.P.G. conducted co-immunoprecipitation analysis with supervision from L.B. J.A., M.C., K.R.D., and K.L.M. performed histopathological analyses and quantification. J.A., and D.M.F. interpreted all datasets. J.A. drafted portions of the manuscript. D.M.F conceived and designed the project, and wrote the manuscript with editorial help from J.A. and A.C.G.

## Methods

### Animal studies and treatment

Animal studies were performed under strict compliance with Institutional Animal Care and Use Committee at University of Pennsylvania (804774). *Kras^LSL-G12D^* [59], *Trp53^flox/flox^*[60], *Trp53^XTR/XTR^*[21], *Rb1^flox/flox^*, *p130^flox/flox^* (*p130* is also known as *Rbl2*) [22] and *Rosa26^FlpO-ER^* mice [61] have previously been described. Mice are mixed B6J/129S4vJae. Mice were transduced with 1.0 × 10^8^ plaque-forming units (PFUs) of Ad:CMV-Cre obtained from University of Iowa Viral Vector Core [62]. For *Trp53* restoration, mice were treated with tamoxifen on two consecutive days with 200μl of a 20 mg ml^−1^ solution dissolved in 90% sterile corn oil and 10% ethanol by oral gavage. Weekly treatment with tamoxifen was administered for long-term experiments. For cyclophilin inhibition, mice were treated with Cyclosporine A with 200μl of a 15mg/kg solution dissolved in 90% sterile corn oil and 10% DMSO. The acquisition of micro-computed tomography (μCT) was performed on μCT setup (MI Labs) with Acquisition 11.0 software. Image reconstruction and visualization was performed with MI Labs REC-11.01. Individual and total tumor volumes were determined by constructing three-dimensional tomograms within ITK-SNAP v3.8 (*itksnap.org*) [63]. Mice in survival studies were monitored for lethargy, labored breathing and weight loss, at which time animals were euthanized.No statistical methods were used to predetermine sample sizes. The size of each animal cohort was determined by estimating biologically relevant effect sizes between control and treated groups and then using the minimum number of animals that could reveal statistical significance using the indicated tests of significance. All animal studies were randomized in ‘control or ‘treated groups. However, all animals housed within the same cage were generally placed within the same treatment group. For histopathological assessments of necrosis, fibrosis, and senescence, researchers were blinded to sample identity and group. Animal sex was randomized at the time of treatment group assignment.

### Allograft studies

SCLC cells were subcutaneously injected into the flanks of NCR^Nude^ mice (Taconic). Tumor volumes were estimated using Vernier calipers and animals were stratified into control or tamoxifen treatment groups when tumors reached a size of ∼100mm^3^. Tumor volumes were recorded bi-weekly through duration of experiment and never allowed to reach a combined tumor volume of 1cm^3^.

### Immunohistochemistry and immunofluorescence

Lung and tumor tissues were dissected into 10% neutral-buffered formalin overnight at room temperature before dehydration in a graded alcohol series. Paraffin-embedded, H&E, and/or Trichrome stained histological sections were produced by the Penn Molecular Pathology and Imaging Core. Immunostaining for cleaved Caspase 3 (1:200,Cell Signaling 9661), ASCL1 (1:200, BD Biosciences 556604), UCHL1 (1:200, Sigma Aldrich HPA005993), CGRP (1:200, Sigma Aldrich C8198), F4/80 (1:500, Novus Biological NB600-404), GFP(1:1000, Abcam 13970), and Ki67 (1:1000, Vector VP-RM04) were performed after citrate-based antigen retrieval. Cleaved Caspase 3 and ASCL1 staining was assessed by immunohistochemistry using ABC reagent (Vector Laboratories, PK-4001) and ImmPACT DAB (Vector Laboratories, SK-4105) according to manufacturer instructions. GFP and Ki67 staining was assessed by immunofluorescence using a biotin-conjugated secondary (Vector), anti-chicken-Alexa594 (Jackson Immunoresearch), and a streptavidin-conjugated Alexa488 antibody.

Immunohistochemistry and immunofluorescence were both performed on paraffin-embedded sections following the same antigen-retrieval protocol. Sections were incubated in primary antibody overnight at 4 °C, secondary antibody for 1 hour at room temperature, and for immunofluorescence Streptavidin-conjugated fluorophore for 1 hour at room temperature in the dark.

For TUNEL staining, tissues were deparaffinized and then permeabilized with 0.1% sodium citrate and 0.1% Triton-X in PBS for 8 minutes. TMR Red-conjugated TUNEL labeling mix (Millipore Sigma, 12156792910) was added to permeabilized tissue sections and incubated for 1 hour at 37 °C in the dark. For all immunofluorescence staining, nuclei were stained using 5 mg/ml DAPI at a 1:1000 dilution for 10 minutes, and then slides were mounted with Fluoro-Gel (EMS, 17985-50). All photomicrographs were captured on a Leica DMI6000B inverted light and fluorescence microscope, or a Leica M80 is a compact stereomicroscope.

### Senescence-associated beta galactosidase (SA-βGal) Staining

SA-βGal staining conducted as previously described at pH 5.5 for mouse cells and tissue [64]. Frozen sections of lung tissue (14μM) or adherent cells were fixed with 0.5% glutaraldehyde in PBS for 15 minutes washed with PBS supplemented with 1mM MgCl^2^ and stained for at least 8hrs at 37°C in PBS containing 1mM MgCl^2^, 1mg/ml X-Gal, 5mM potassium ferricyanide and 5mM potassium ferrocyanide. Tissue sections were counterstained with eosin.

### Histological quantification

ImageJ software was used to determine the frequency of cells that were positive for specified antigens (Ki67) as a fraction of total tumor cells. Data points represent individual tumors for Ki67 staining. For SA-βGal and TUNEL a binary staining score for each tumor was determined based on staining intensity scored from 0 (minimal staining) to 1 (ubiquitous staining). For Trichrome staining, a similar staining scoring system was applied where individual tumors were scored from 0 (minimal necrosis or fibrosis), to 1 (ubiquitous necrosis or fibrosis). PennVet Comparative Pathology Core developed a system to stratify SCLC tumors based on immune infiltration scored from 0 (no neutrophil or mononuclear infiltration) to 4 (high neutrophil or mononuclear infiltration). Data points represent either individual animals or tumors, as indicated.

### Cell Lines

Primary mouse tumors were mechanically separated using scissors under sterile conditions and cultured in high-glucose DMEM supplemented with 10% fetal bovine serum, GlutaMAX and antibiotics at 37°C and 5% CO_2_ until cell line establishment. Cell lines were authenticated for genotype. The p53^XTR/XTR^ allele was validated via three-primer PCR reactions using purified genomic DNA as a template to detect *XTR*, *TR*, *R* and *WT* alleles. Primers used were as follows: (1) 5′-cttggagacatagccacactg −3′, (2) 5′-caactgttctacctcaagagcc −3′ and (3) 5′-cttgaagaagatggtgcg −3′. Cell lines were further validated for the expected, genotype-associated protein expression patterns by western blot. KP^Restorable^ cell lines used in senescence experiments were derived from *Kras^LA2/+^;Trp53^LSL/LSL^;Rosa26^CreERT2/CreERT2^*adenocarcinomas as previously described (Feldser, et. al. 2010). All cell lines were tested for mycoplasma using MycoAlert detection kits (Lonza) as per manufacturer instructions.

Cell number was counted using a Beckman Coulter Z2 Cell and Particle Counter. 4-Hydroxytamoxifen (4-OHT) dissolved in ethanol was administered once at the time of cell plating at a final concentration of 500 nM. AdCre or AdFlpO was purchased from the University of Iowa Gene Vector Core and administered to adherent cells at time of cell plating. Proliferation of cell lines was determined by plating the indicated cell number and analysed by Coulter Counter cell counts at the indicated time points. Photomicrographs were captured on a Leica DM IL LED inverted light and fluorescence microscope. For live cell imaging, photomicrographs were obtained in a 37°C humidified chamber every 15 min for 4 days using a Leica DMI6000B microscope.

HEK293FT cells that were used for lentivirus production were obtained from Invitrogen. HEK293FT cells were maintained in DMEM containing 10% Bovine Serum. HEK293FT cells were validated by verifying that high-titer virus production was possible. Lentivirus production was performed as described previously [65].

### Immunoblot analysis

Cells were lysed in RIPA buffer, resolved on NuPage 4–12% Bis-Tris protein gels (Thermo Fisher) and transferred to polyvinylidene fluoride (PVDF) membranes. Blocking, primary and secondary antibody incubations were performed in Tris-buffered saline (TBS) with 0.1% Tween-20. p53 (1:2500, Leica Biosystems, P53-PROTEIN-CM5), p21 (1:500, Santa Cruz, sc-6246), Cleaved Caspase-8 (1:1000,Cell Signaling Technology, 8592S), Cleaved Caspase-3 (1:1000,Cell Signaling Technology, 9661S), Cleaved PARP (1:1000,Cell Signaling Technology, 9548S), RB (1:1000, Abcam, ab181616), ASCL1 (1:1000, BD Biosciences 556604), PPIF (Cyclophilin D, 1:1000, Abcam, ab110324), PPIA (Cyclophilin A, 1:1000, Cell Signaling Technology, 2175S), β-actin (1:10000,Sigma Aldrich, A2066), HSP90 (1:10000, BD Transduction Laboratories, 610418), GAPDH (1:10000,Cell Signaling Technology, 2118S), COXV (1:4000,Cell Signaling Technology, 11967S), H3 (1:10000, Abcam ab1791) were assessed by western blotting. HSP90, GAPDH, β-actin, COXIV, and H3 were used as loading controls. Protein concentration was determined using a BCA protein assay kit (Pierce).

### Flow Cytometry

For *in vitro* experiments, single cell suspensions were prepared from SCLC cells after collection by passing specimens through a 100-μM cell strainer. Cells were resuspended in FACS buffer. Equal cell numbers of cells were plated and stained for flow cytometry analysis. Cell cycle analysis was performed according to the manufacturer’s instructions for the APC BrdU Flow Kit (BD Pharmingen). TUNEL staining was performed according to the manufacturer’s instructions for the In Situ Cell Death Detection Kit, TMR red (Millipore Sigma, 12156792910). For cell viability assays, cells were resuspended in FACS buffer containing DAPI at a 1:1000 dilution. For all *in vitro* experiments, flow cytometry was performed using an Attune NxT flow cytometer (Thermo Fisher). Data were analyzed using FlowJo v10.8. Data points represent technical replicates, or means from independent experiments, as indicated.

For *in vivo* experiments, tumors were microdissected directly from the lungs of *RP^X^R2* mice and individually placed in 500 μl of tumor digestion buffer consisting of PBS containing 10 mM HEPES pH 7.4, 150 mM NaCl, 5 mM KCl, 1 mM MgCl_2_, and 1.8 mM CaCl_2_, along with freshly added Collagenase 4 (Worthington 100 mg/ml solution, 20 μl per ml of digestion buffer) and DNase I (Roche 10 mg/ml solution, 4 μl per ml of digestion buffer). Tumors were manually disassociated using scissors, and then placed in a 4 °C shaker for 1 hour at 250 rpm. Digested tumors were then filtered into strainer-cap flow tubes (Corning, 352235) containing 1 ml of horse serum (Thermo Fisher, 16050122) to quench the digestion reaction. Cells were spun down at 200 g for 5 minutes with the cap in place to obtain all cells. The supernatant was aspirated, cells were washed once with PBS and resuspended in FACS buffer. Cells were labeled with the following antibodies: PD-1 FITC (1:50, Biolegend), NKp46-PE (1:50, Biolegend), CD103-PETR (1:100, Biolegend), CD3-PE/Cy5 (1:100, Biolegend), CD8-PE/Cy7 (1:100, Biolegend), CD44-APC (1:100, Biolegend), CD45-AF700 (1:400, Biolegend), F4/80-APC/Cy7 (1:100, Biolegend), CD11b-PerCP/Cy5.5 (1:200, BD Biosciences), CD11c-QD605 (1:100, Biolegend), CD45-BV570 (1:100, Biolegend), CD4-BV650 (1:50, Biolegend), Live/Dead Aqua (1:600, Invitrogen). For all *in vivo* experiments, flow cytometry was performed using a BD LSR II flow cytometer (BD Biosciences). Data were analyzed using FlowJo v10.8. Data points represent individual tumors.

### Drugs, cell death inhibitors

Drugs or cell death inhibitors are used at the following concentrations: Necrostatin-1s (10μg/mL, Selleck Chemicals), Ferrostatin-1 (2μM, XcessBio), Liproxstatin-1 (2μM, Sellek Chemicals), Vitamin E (125μM, Sigma Aldrich), Cyclosporin A (1μM, Sigma Aldrich), ZVAD-FMK (20μM, Abcam), NIM-811 (1μM, MedChem Express), FK506 (1μM, ApexBio). Cell death inhibitors were added to cells at time of plating for the duration of the experiment.

### CRISPR design and production

A validated sgRNA targeting *Atg5* was obtained from Genscript. All remaining sgRNAs used were designed using the Sanjana Lab CRISPR Cas9 library design tool (http://guides.sanjanalab.org). A minimum of 2 guides were designed per target. sgRNAs targeting *βgal* and *Trp53* were used as experimental controls [66]. All sgRNA duplexes were Golden Gate-cloned into BsmBI sites of the LentiCRISPRv2-Puro or LentiCRISPRv2-mCherry vectors [65, 67, 68]. The sgRNA sequences used for targeting Cas9 are detailed in Supplementary Table 1.

### Transfection-mediated gene transfer

HEK293T cells were transfected with plasmids using polyethylenimine (PEI) (Polysciences, #24765). 4.0μg total plasmid DNA was transfected for 48 h.

### Immunoprecipitation and immunoblot

Cells were lysed in NP-40 buffer (0.1% NP-40, 15mM Tris–HCl pH7.4, 1 mM EDTA, 150 mM NaCl, 1 mM MgCl2, 10% Glycerol) containing protease inhibitors (Sigma, #11697498001) and the lysates were incubated with anti-FLAG M2 Affinity Gel (Sigma, #A2220) at 4°C for 2 h. After five washes with NP-40 buffer, the anti-FLAG Gel was mixed with Laemmli buffer and boiled at 95°C for 5 min. After SDS–PAGE electrophoresis and transfer, primary antibodies and HRP-linked secondary antibodies were incubated with the membrane for 2 h at room temperature and overnight at 4°C, respectively. After washing with PBS-T three times and PBS once, the membrane was detected by the chemiluminescence system (Thermo Fisher Scientific, #32106). Where indicated, Cyclosporin-A was included during lysis, anti-FLAG Gel incubation, and subsequent washes. The following antibodies were used: anti-FLAG (Sigma, #F7425, 1:6,000), anti-HA (Cell Signaling Technology, #3724, 1:2,000), and anti-Rabbit IgG-HRP (GE Healthcare, #NA934V, 1:10,000).

### Plasmids

pcDNA3.1+C-HA Trp53 (OMu22847) and pcDNA3.1+C-FLAG *Ppia* (OMu14516) plasmids used in mammalian over-expression experiments were purchased from Genscript. The detailed information regarding plasmid constructions is available by request. Briefly, cDNA-expression was achieved by intramolecular ligation cloning strategies into the pcDNA3.1 transient mammalian expression vector. Domain deletions in the mouse *Trp53* cDNA were generated using the following targeted PCR-amplification primers:

BamHI_ATG_*mTrp53*-WT_forward: cgcggatccATGACTGCCATGGAGGAGTCACAG

BamHI_ATG_*mTrp53*-ΔTAD_forward: cgcggatccATGCTCCGAGTGTCAGGAGCTCC

BamHI_ATG_*mTrp53*_ΔTAD,Proline_forward:

cgcggatccATGCTGTCATCTTTTGTCCCTTCTCAAAAAACTTAC

BamHI_ATG_*mTrp53*-ΔTAD,Proline,SH3/DBD_forward:

cgcggatccATGCCAGGGAGCGCAAAGAGAGC

EcoRI_no-STOP_*mTrp53*-WT_reverse: agtgaattcGTCTGAGTCAGGCCCCACTTTC

EcoRI_no-STOP_*mTrp53*-ΔCterm_reverse: agtgaattcGGGCAGTTCAGGGCAAAGGAC

### Seahorse XF Cell Mito Stress Analysis

Oxidative respiration was measured using XF Cell Mito Stress Test Kit (Agilent Technologies, 103015-100). 1 x 10^4^ cells per well were seeded on an XF96 Cell Culture Microplate. Each cell line was split into two groups: control (EtOH) or restored (4-OHT) and 6 technical replicates were plated. Microplate was incubated for 24h at 37C. Seahorse XF96 FluxPak sensor cartridge was hydrated with 200 μl of Seahorse Calibrant in a non-CO2 incubator at 37C overnight. After 24h, cells were incubated with base medium (Agilent Technologies, 102353-100) containing 2 mM L-glutamine, 1 mM sodium pyruvate, and 10 mM glucose in a non-CO2 incubator at 37C for 45 min prior to assay. Oxygen consumption rate (OCR) was measured by XFe96 extracellular flux analyzer with sequential injections of 1 μM oligomycin, 1 μM FCCP, and 0.5 μM rotenone/antimycin A. After the run, cells were lysed with 15 μl RIPA buffer and protein concentration was quantified using BCA Protein Assay Kit (Thermo Scientific). OCR measurements were normalized to the protein concentration in each well.

### Cellular Protein Fractionation

Cells were collected and lysed in hypotonic buffer (20mM Tris pH 7.5, 5mM MgCl_2_, 5mM CaCl_2_, 1mM DTT, 1mM EDTA, protease inhibitor) for 30 minutes on ice. Cells were dounced homogenized and spun down at 3000rpm for 20min. Cytosolic fraction was obtained by collecting the supernatant. Nuclear fraction was obtained by resuspending nuclear pellet in 1mL low salt buffer (20mM Tris pH 7.5, 5mM MgCl_2_, 20mM KCl, 5mM CaCl_2_, 1mM DTT, 1mM EDTA), adding 1mL (100uL x10) of high salt buffer (20mM Tris pH 7.5, 5mM MgCl_2_, 1.2M KCl, 5mM CaCl_2_, 1mM DTT, 1mM EDTA) with mixing every 100uL, and incubating the mixture at 4°C for 2h with gentle agitation. Mixture was spun down at 25k g for 15 minutes, and supernatant (nuclear fraction) was collected. All fractions were TCA precipitated, pellets were resuspended in denaturing buffer (50mM Tris pH 8.3, 5mM EDTA, 0.05% SDS, and 6M urea), and protein concentration was quantified using BCA Protein Assay Kit (Thermo Scientific). For mitochondrial fractionations, cells were processed using Pierce Mitochondria Isolation Kit for Cultured Cells (Thermo Scientific) as per the manufacturer’s instructions.

### RNA Isolation and RNA-sequencing

3 Type D and 3 Type V SCLC tumor-derived cell lines were treated with either vehicle (EtOH) or 4-OHT for 3 days and RNA was isolated using the RNeasy Mini Kit (Qiagen) as per the manufacturer’s instructions. RNA concentrations were measured using the Qubit RNA Assay Kit (Invitrogen) and sample quality was determined using the RNA 6000 Nano Kit on a 2100 BioAnalyzer (Agilent). Sequencing libraries were prepared using the TruSeq Stranded mRNA Library Prep Kit (Illumina) as per the manufacturer’s instructions and library quality was determined used the Agilent DNA 1000 Kit on a 2100 BioAnalyzer. Validated libraries were subjected to 75-bp single-end sequencing on the Illumina NextSeq 500 platform. Fastq files for each sample were aligned against the mouse genome mm10 (GRCm39) using the Kallisto pseudoaligner, or HISAT2. Differential expression analysis was carried out with the Limma tool in R, or DeSeq2. For volcano plot data, differentially enriched genes were defined as genes with a false-discovery rate adjusted *P* value less than 0.05 and log_2_-normalized fold change in expression greater than ± 1. Geneset enrichment analysis was done using the GSEABase and msigdbr packages in R, or using the GSEA v4.2.2 software provided by the Broad institute (https://www.gsea-msigdb.org/gsea/index.jsp). ‘Type D’ and ‘Type V’ gene signatures were defined as genes with a false-discovery rate adjusted *P* value less than 0.05 and log_2_-normalized fold change in expression greater than 1 (109 genes were differentially regulated in Type D cells, 57 genes were differentially regulated in Type V cells). Human SCLC datasets for SCLC-ASCL1 and SCLC-NEUROD1 signature enrichment were obtained from CellMiner-SCLC (https://discover.nci.nih.gov/SclcCellMinerCDB/) [46]. Single sample GSEA was performed using the GSVA package in R. Gene ontology analysis was performed using the gProfiler2 package in R, or the g:Profiler web tool (https://biit.cs.ut.ee/gprofiler/gost). Visualization of gene expression patterns (heatmaps) and GSEA data was performed using the gplots package in R. Other visualization and statistical analysis were performed using Prism 9.

### Chromatin Immunoprecipitation (ChIP)

3 Type D and 3 Type V SCLC tumor-derived cell lines were treated with either vehicle (EtOH) or 4-OHT for 2 days for ChIP. 1 x 10^7^ SCLC cells were harvested and cross-linked at room temperature by resuspending in PBS containing 1% formaldehyde for 2.5 minutes. The reaction was quenched by addition of glycine to a final concentration of 0.125 M. After washing with cold 1X PBS, cells were lysed in 1mL sonication buffer (0.25% Sarkosyl, 1 mM DTT, and protease inhibitors into RIPA buffer). Cell suspension was sonicated using a Covaris S220 Focused Ultrasonicator until chromatin was sheared to a size range of around 200bp. Lysates were centrifuged 14k rpm for 5 minutes and supernatant was collected. After saving 10% for an input sample, 20uL of anti-p53 antibody (CM5, Leica Biosystems was added to lysate and incubated overnight. 200uL of a bead slurry prepared from Protein A/G Magnetic Beads (Thermo Scientific) were added to lysates and incubated in a rotator at 4°C for 4h. Chromatin-bound beads were washed three times with low-salt wash buffer (0.1% SDS, 1% Triton X-100, 1 mM EDTA pH 8.0, 50 mM Tris-HCl at pH 8.0, 150 mM NaCl), high-salt wash buffer (0.1% SDS, 1% Triton X-100, 1 mM EDTA pH 8.0, 50 mM Tris-HCl pH 8.0, 500 mM NaCl), and LiCl wash buffer (150mM LiCl, 0.1% SDS, 0.5% deoxycholic acid sodium salt, 1% NP-40, 1 mM EDTA pH 8.0, 50 mM Tris pH 8.0). After washing the beads once with TE buffer containing 50mM NaCl, the beads were resuspended in ChIP elution buffer (1% SDS, 200 mM NaCl, 10 mM EDTA pH 8.0, 50 mM Tris pH 8.0) and eluted for 30 min at 65°C. Bead solution was spun down for 1 min at 16g and 200uL of the supernatant were transferred to a new tube. 10% input was diluted to 200uL using ChIP elution buffer. Diluted input and experimental samples were reverse cross-linked overnight at 65°C. All samples were treated with RNAse A and incubated at 37°C for an hour, followed by Proteinase K treatment and incubated for an hour at 55°C. DNA was recovered by PCR Purification Kit (QIAGEN) and eluted into 50uL of molecular biology grade water. Chromatin-immunoprecipated DNA was validated for p53 target enrichment by qPCR using SYBR Green Power Up (Thermo Fisher) and ViiA7 Real-Time PCR System (Thermo Fisher). Sequencing libraries were prepared using the NEBNext Ultra II DNA Library Prep kit for Illumina (New England Biolabs) size selection, as per the manufacturer’s instructions, with the following conditions: 30uL ChIP sample or 5uL input sample used when end prepping, adaptor was diluted 10-fold for ChIP samples but not diluted for input, 200 bp bead based size selection used for all samples, 14 cycles used for PCR enrichment of ChIP samples and 7 cycles for input samples. Library quality was determined used the Agilent DNA 1000 Kit on a 2100 BioAnalyzer (Agilent). Validated libraries were subjected to 75-bp single-end sequencing on the Illumina NextSeq 500 platform.

### ChIP-sequencing

p53 ChIP-seq data was aligned to mouse genome mm10 (GRCm39) by bowtie2 with default parameters. Peak calling was performed by MACS2 with a p-value threshold of 1E-10. ChIP-seq signal in a specific region was calculated by the normalized RPM. Briefly, ChIP-seq reads aligning to the region were extended by 100 bp and the density of reads per bp was calculated using featureCounts. The density of reads in each region was normalized to the total number of million mapped reads, producing read density in units of reads per million mapped reads per bp (RPM per bp). Differential p53 binding analysis between Type D and Type V cells was carried out using the DiffBind tool in R. Differential regions were identified with fold change and p-value cutoffs, as indicated. The nearest gene of specific p53 peak was treated as the associated gene of p53, which was identified by the HOMER module annotatePeaks.pl.

### Statistical analysis

All analyses were performed in the Graphpad Prism 9 software package. For survival studies, log-rank (Mantel–Cox) tests were performed to determine significance. For SA-βGal contingency analysis, Fisher’s exact test was performed. For cyclophilin CRISPR knockout or inhibition, and ferroptosis inhibition experiments, Dunnett’s multiple comparison tests were performed. For all remaining experiments, unpaired Student’s *t-*tests were performed to determine significance, as indicated.

---

**Fig. S1.**
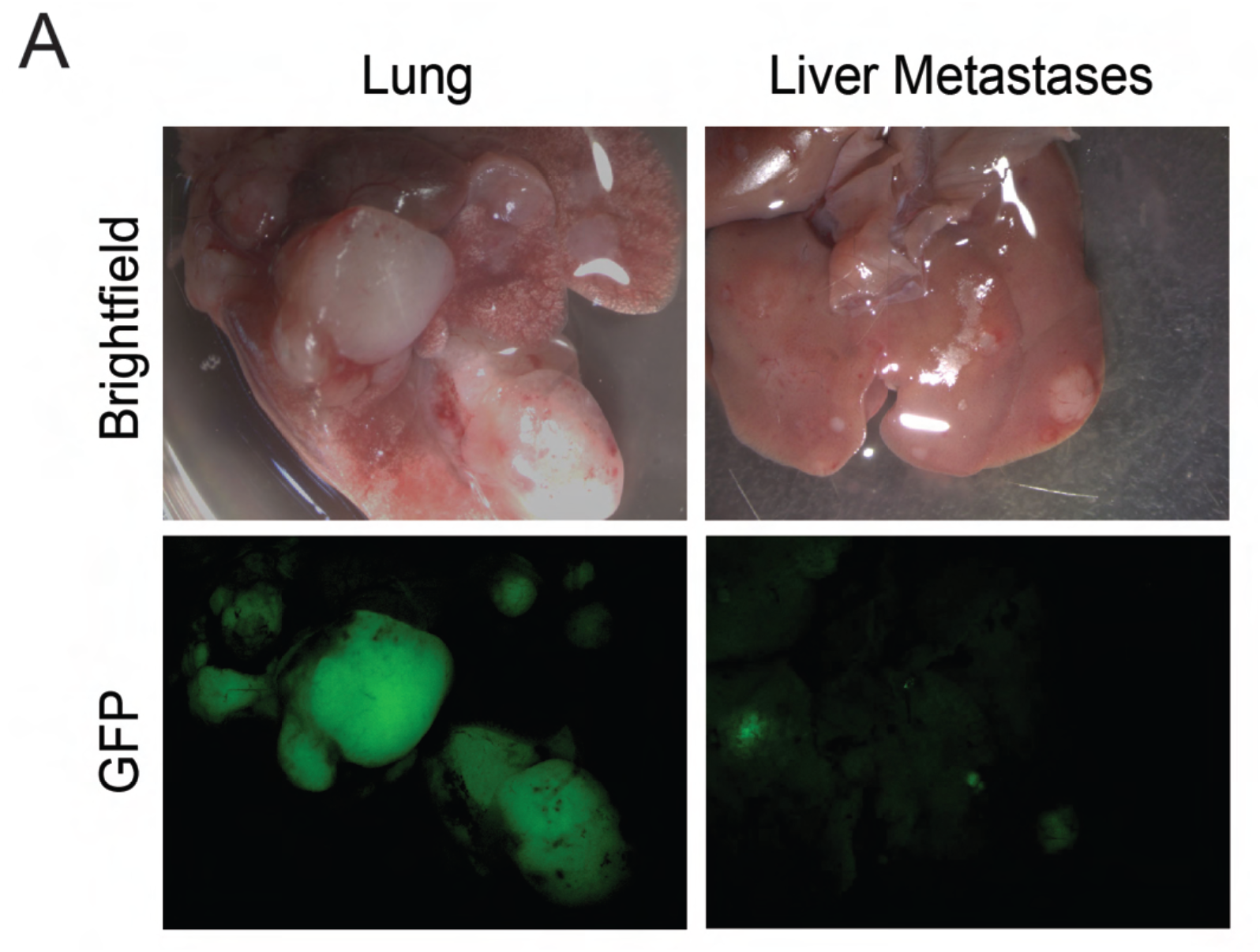
*RP^R^R2* animals succumb to rare *RP^TR^R2* GFP+ cells that escape tamoxifen-induced p53 restoration. **(A)** Representative brightfield and fluorescent micrographs of lungs and liver metastases from *RP^R^R2* mouse treated with tamoxifen for 14 weeks.

**Fig. S2.**
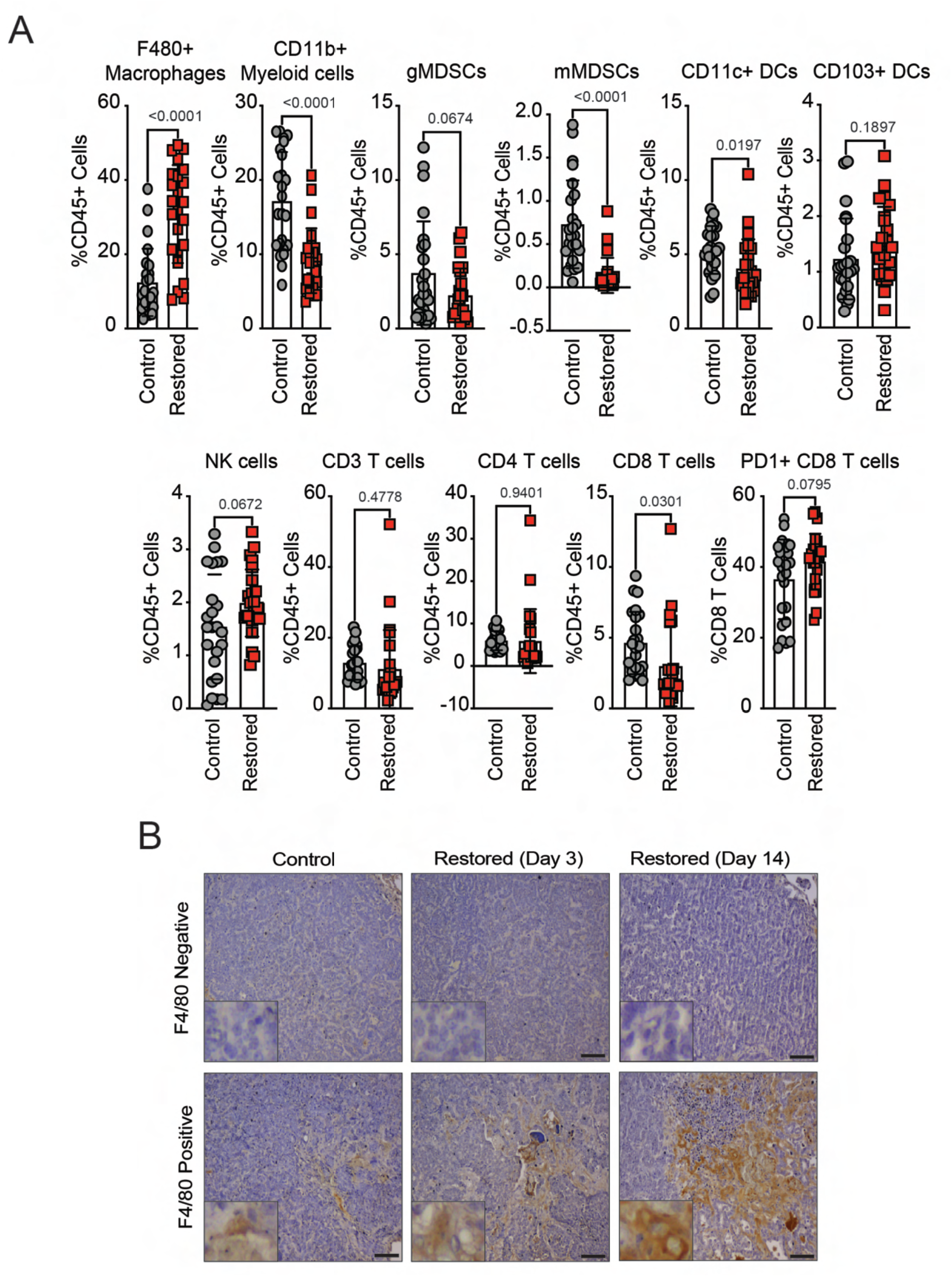
p53 reactivation per se does not induce widespread immune inflammation but necrotic areas are associated with increased macrophage infiltration. (**A**) Flow cytometry analysis of macrophage, myeloid-derived suppressor cells (MDSCs), dendritic cells (DC), natural killer cells (NK), and T-cell infiltration in *RP^TR^R2* tumors 3 days after vehicle (Control) or tamoxifen (Restored) treatment. n=24 tumors from 4 Control mice, n= 24 tumors from 3 Restored mice. Statistical significance was determined by Student’s *t-*test. Error bars represent mean ± s.d. (**B**) Representative photomicrographs for F4/80 IHC in Control or Restored RP^X^R2 tumors. Scale bars, 25µm: insets are magnified 5×.

**Fig. S3.**
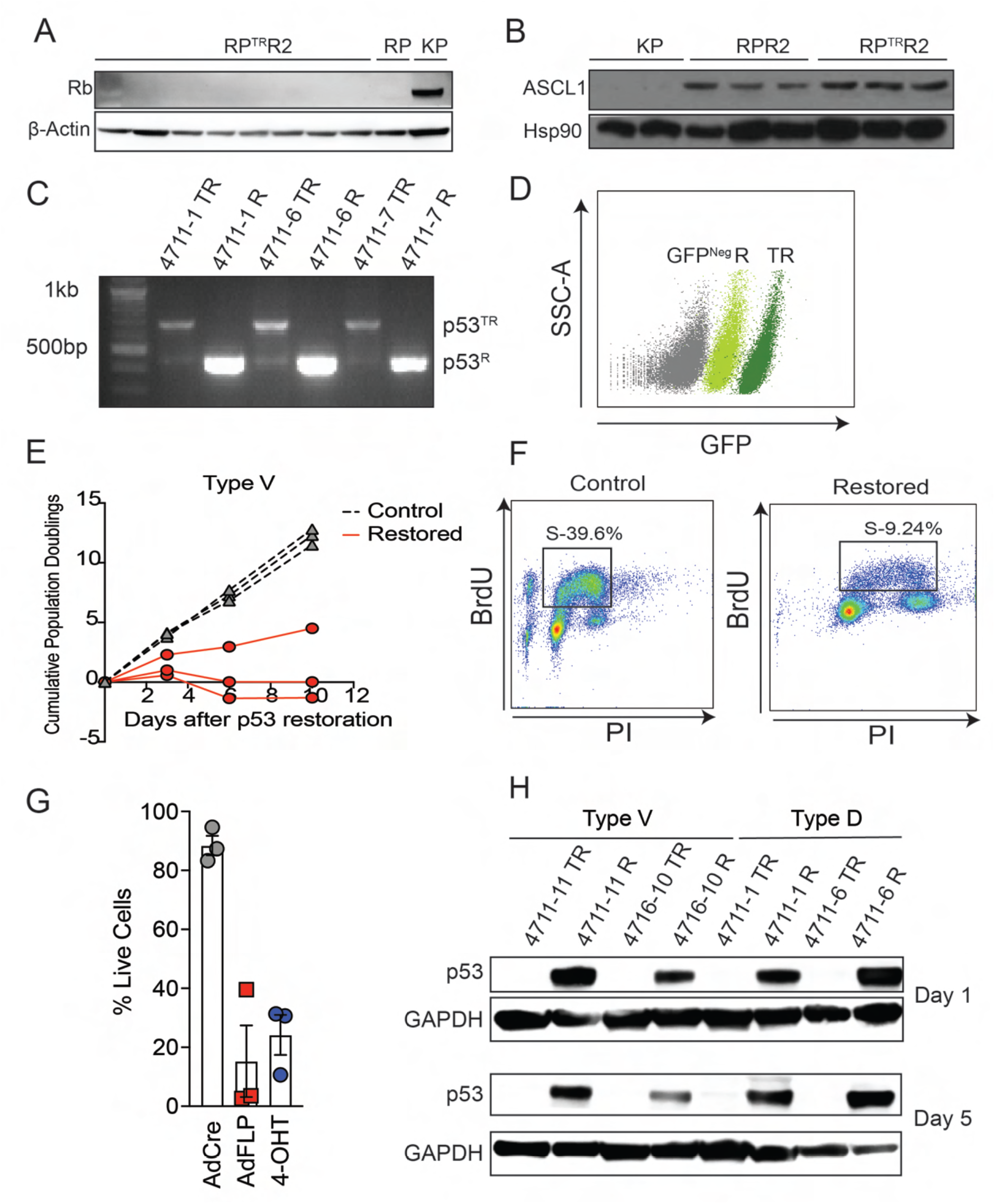
p53 reactivation induces death or senescence in tumor-derived SCLC cell lines. (**A**) Immunoblot analysis for Rb in *RP^TR^R2* tumor-derived cell lines. *RP* and *KP* cell lines used as negative and positive controls for Rb expression, respectively. β-actin used as loading control. (**B**) Immunoblot analysis for expression of the neuroendocrine marker, ASCL1, in *RP^TR^R2* tumor-derived cell lines. *KP* and *RPR2* cell lines used as negative and positive controls for ASCL1 expression, respectively. Hsp90 used as loading control. (**C**) PCR-based detection of *p53^TR^* and *p53^R^* alleles in *RP^TR^R2* cell lines (n=3) after 24hrs of 4-OHT treatment. (**D**) Detection of GFP reporter expression from *TR* alleles in *RP^TR^R2* cell lines by flow cytometry analysis. *KP* cell line used as GFP^neg^ control. Representative of n=8 cell lines. (**E**) Cumulative population doublings of Type V cells in Fig. 3E. 150,000 cells plated for n=3 Type V cell lines at t=0 hrs. 150,000 cells cells were re-plated at t=3 days and t=6 days. (**F**) Flow cytometry plots of BrdU incorporation in Type V cells in (**3f**). Percentage of cells in S-phase (BrdU+) was calculated in Type V cells (n=3) 72hrs after 4-OHT treatment. (**G**) Flow cytometry-assisted cell viability assay in Type D cells 4 days after 4-OHT treatment, or Ad:FlpO treatment. Ad:CMV-Cre treatment used as a negative control for p53 reactivation. Each symbol represents the mean of n=3 technical replicates from a Type D cell line. Distinct Type D cell lines (n=3) were used as biological replicates. Error bars represent mean ± SEM. (**H**) Immunoblot analysis for expression of p53 in Type V and Type D cell lines one or five days after 4-OHT treatment. n=2 representative cell lines used per cell line. GAPDH used as loading control.

**Fig. S4.**
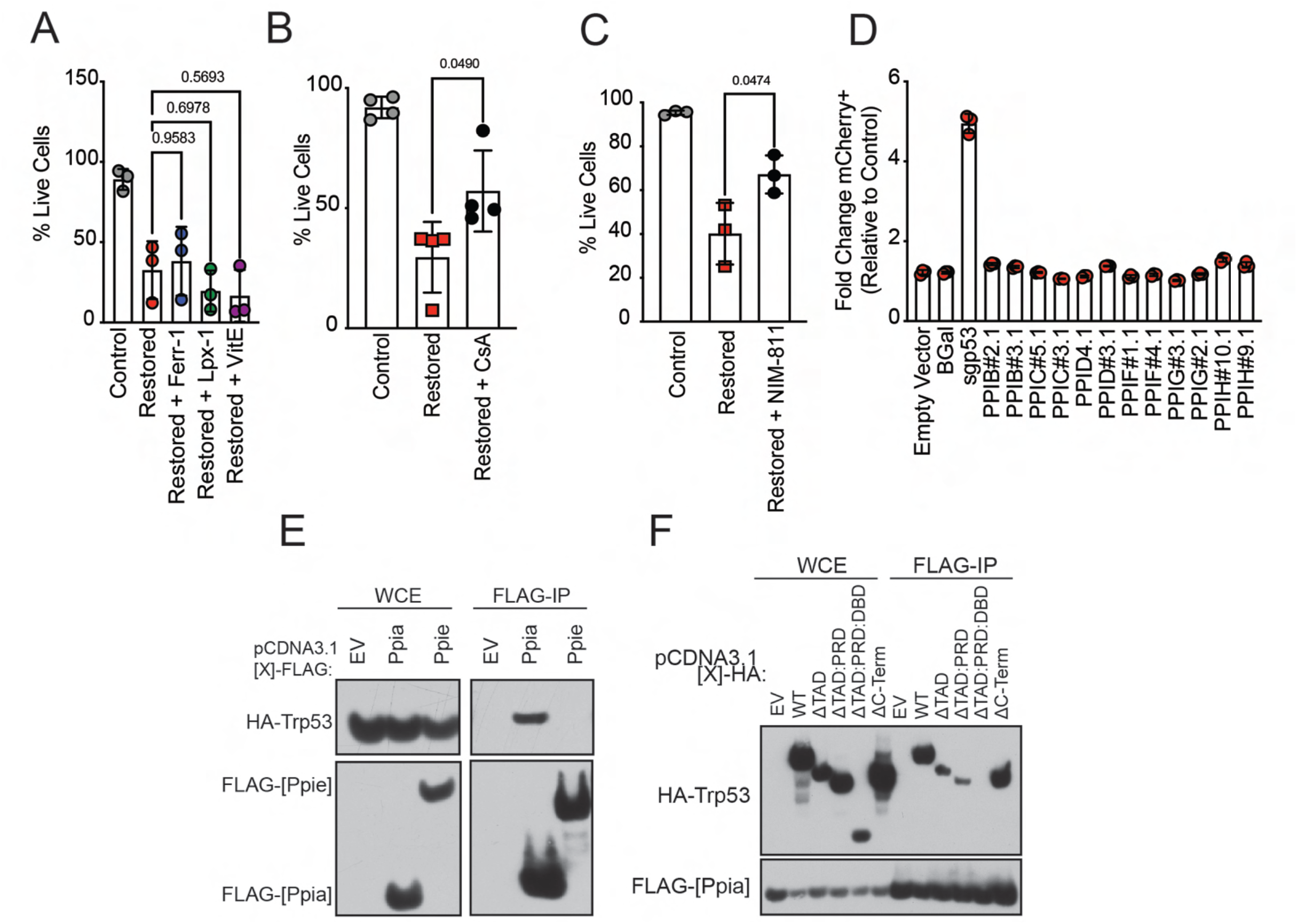
p53 induces Cyclophilin-dependent cell death in SCLC. (**A**) Type D cells were treated with 4-OHT and ferroptosis inhibitors for 4 days. Percentage of live cells was determined using flow cytometry by quantification of DAPI negative population. Each symbol represents the mean of n=3 technical replicates from a Type D cell line. Distinct Type D cell lines (n=3) were used as biological replicates. Statistical significance was determined by Dunnett’s multiple comparison test. Error bars represent mean ± s.d (B) Type D cells were treated with 4-OHT and CsA for 3-5 days. Percentage of live cells was determined using flow cytometry by quantification of DAPI negative population. Each symbol represents the mean of n=3 technical replicates from a Type D cell line. Distinct Type D cell lines (n=4) were used as biological replicates. Statistical significance was determined by Student’s *t*-test. Error bars represent mean ± s.d. (**C**) Type D cells were treated with 4-OHT and NIM-811 for 3 days. Percentage of live cells was determined using flow cytometry by quantification of DAPI negative population. Each symbol represents the mean of n=3 technical replicates from a Type D cell line. Distinct Type D cell lines (n=3) were used as biological replicates. Statistical significance was determined by Student’s *t*-test. Error bars represent mean ± s.d. (**D**) Quantification of fold change mCherry+ cells after 72hrs of 4-OHT treatment in additional Type D cell line expressing sgRNAs targeting distinct cyclophilins. Empty Vector, β-Gal, and a p53 targeting sgRNA used as controls. Each symbol represents a technical replicate (n=3). Error bars represent mean ± s.d. Experiment was conducted in n=2 biological replicates (Figure 3D). (**E**) Immunoblot analysis for PPIA, PPIE and p53 expression from WCE or FLAG-IP samples from HEK293T cells overexpressing FLAG-*Ppia* or FLAG-*Ppie,* and HA-*Trp53.* (**F**) Immunoblot analysis for PPIA and p53 expression from WCE or FLAG-IP samples from HEK293T cells overexpressing FLAG-*Ppia* and distinct truncated mutations of HA-*Trp53.* EV=empty vector; WT= wild-type; TAD= transactivation domain; PRD=proline rich domain; DBD= DNA-binding domain.

**Fig. S5.**
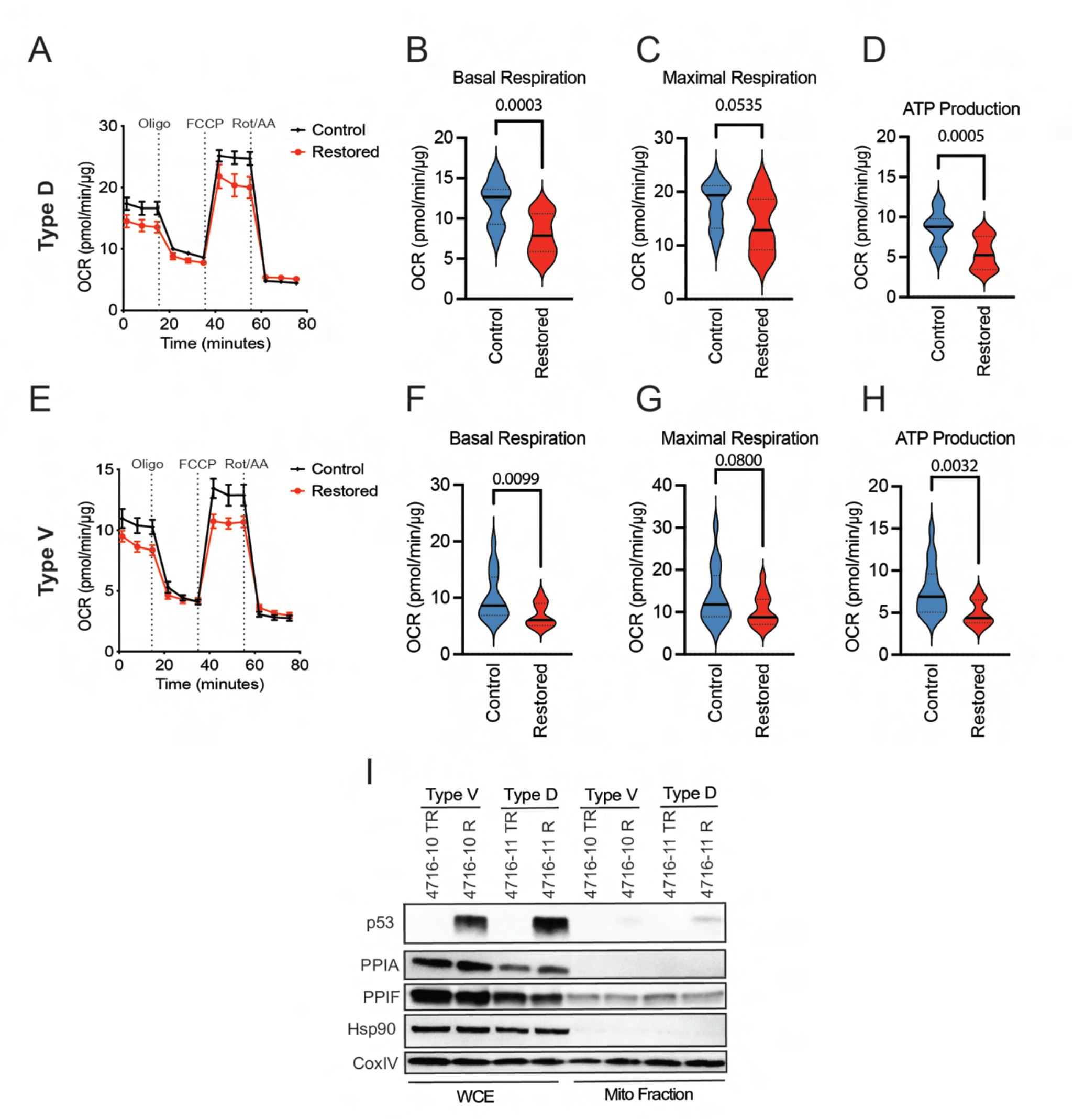
p53 does not prominently associate with the mitochondria and mitochondria are not dysfunctional during Type D SCLC cell death. Seahorse XF cell mitochondrial stress test assay performed in Type D (**A**) and Type V (**E**) cells after treatment with vehicle or 4-OHT. (**A,E**) OCR profile plots. (**B,F**) Basal respiration. (**C,G**) Maximal respiration. (**D,H**) ATP production. Relative oxygen consumption rate was normalized to protein abundance. Each symbol in OCR profile plots represents the mean of at least n=4 technical replicates of three reading cycles. Error bars represent s.d. Experiment was conducted in n=3 cell lines per subtype. (**I**) Immunoblot analysis from a representative Type D or Type V cell line for PPIA, PPIF, and p53 expression in whole cell extracts (WCE) and mitochondrial fractions after 4 days of 4-OHT treatment. Hsp90 used as loading control for WCE; COXIV used as a loading control for mitochondrial fraction.

**Fig. S6.**
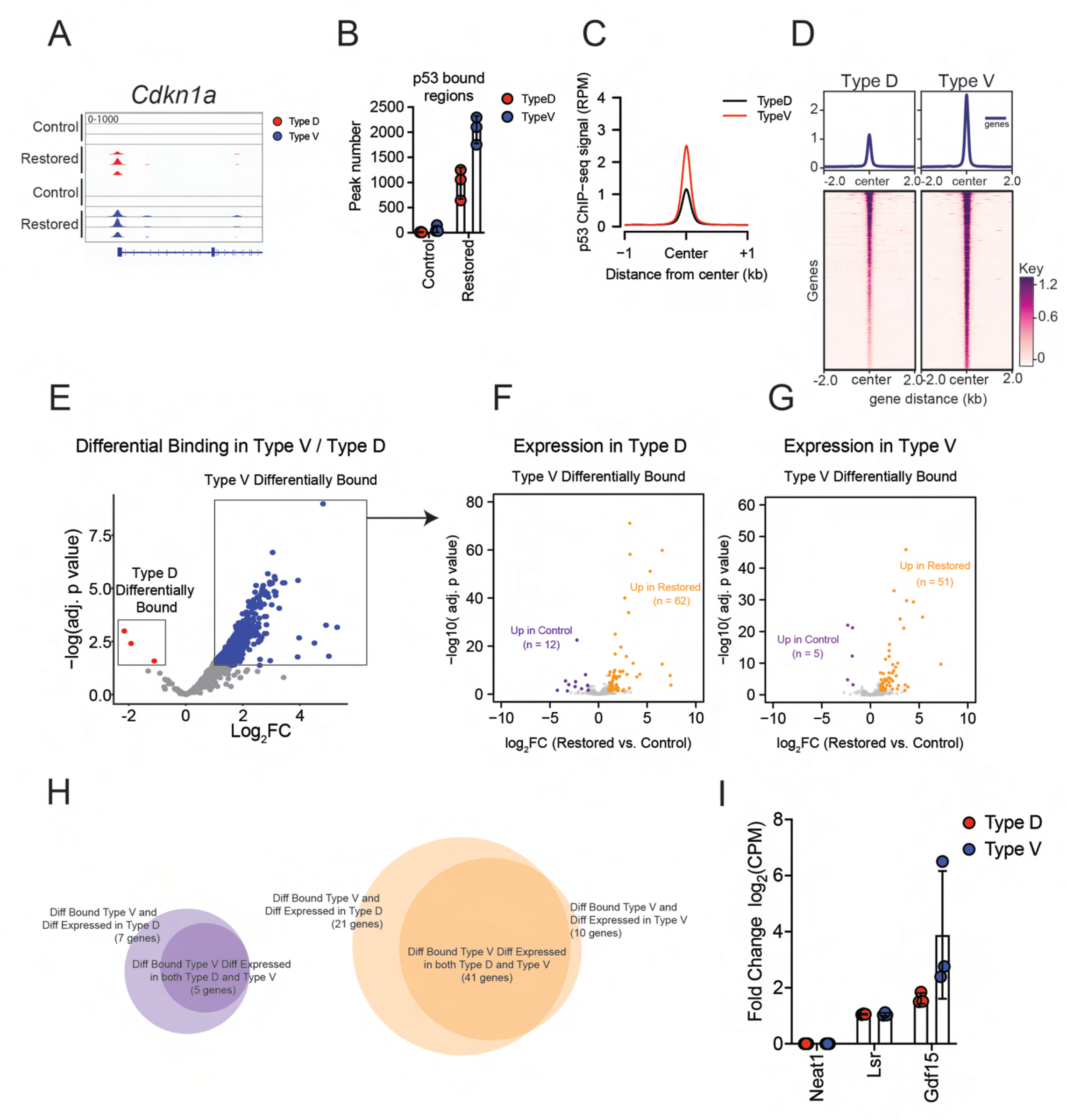
p53 binding does not identify subtype-specific transcriptional programs. (**A**) Genome browser view of p53 ChIP-seq signal at a canonical p53-target gene, *Cdkn1a.* (**B**) Quantification of p53 bound regions in Control and Restored cells. n=3 for both Type D and Type V cells. (**C**) Comparison of p53 ChIP-seq signal between Type D and Type V cells. Data are centered on p53 bound peaks across a ± 1kb window. (**D**) Heatmap representation of p53 bound chromatin peaks 48hrs after 4-OHT treatment in Type D (n=3) and Type V cells (n=3). Heatmaps are centered on p53-bound peaks across a ± 2kb window. (**E**) Volcano plot of differentially p53 bound regions in Type V (n=1032 genomic regions; blue) and Type D (n=3 genomic regions; red) cells. (**F,G**) Volcano plots of RNA-sequencing gene expression from Type D (F) and Type V (G) cells of Type V differentially bound p53 regions identified in (E). Colored dots represent genes that are differentially enriched (log_2_ fold-change greater-than 1 and false discovery rate (FDR)-adjusted *P*-value less-than 0.05) in Restored (orange) or Control (purple). (**H**) Venn diagrams comparing differentially bound Type V genes and their differential expression in Control (purple) or Restored (orange) cells. **(I**) Quantification of log_2_ fold change in counts per million (CPM) of Type D differentially bound genes identified in (F) in Type D (n=3) and Type V (n=3) cells. Error bars represent mean ± s.d.

**Fig. S7.**
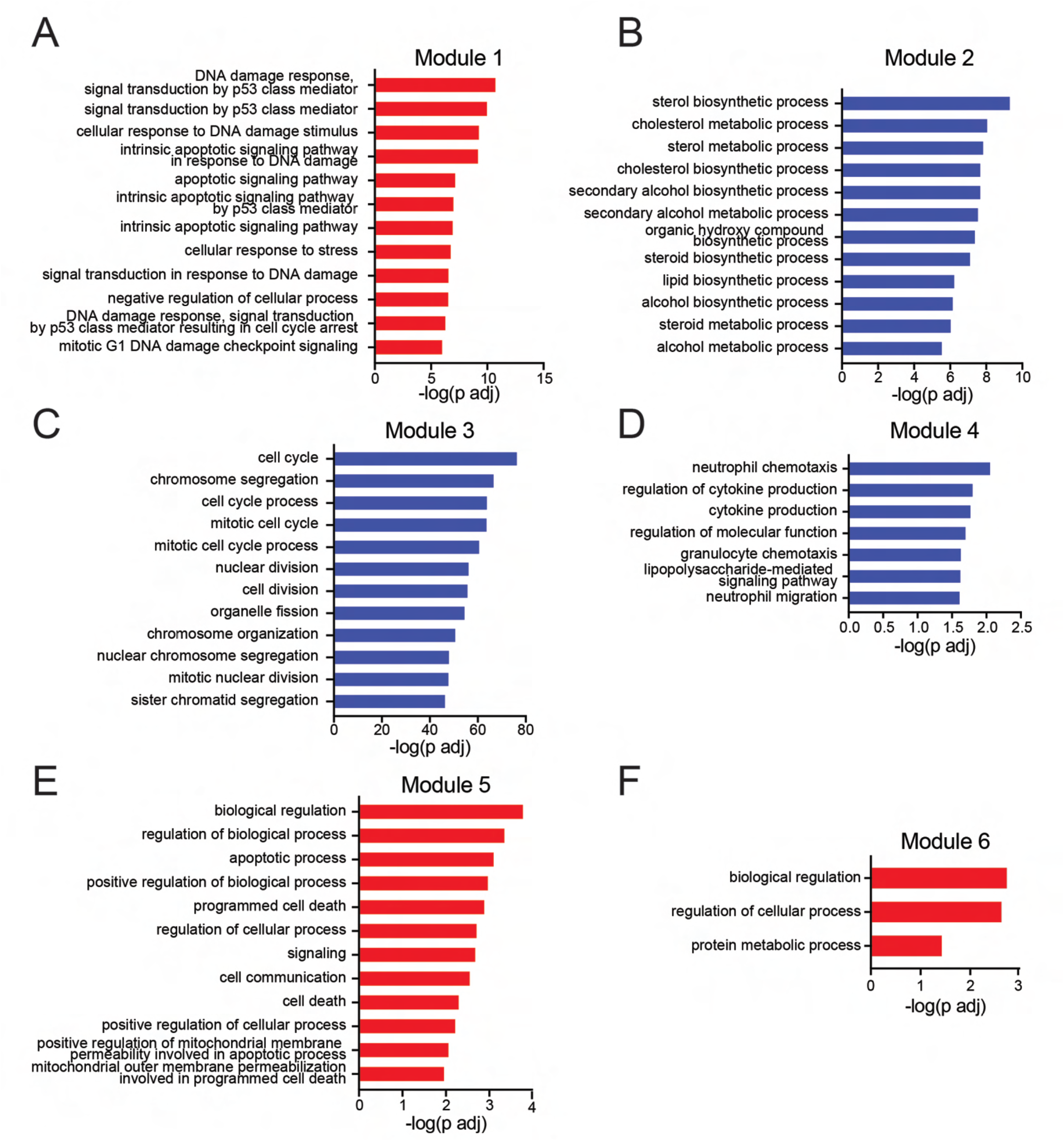
p53 reactivation modulates diverse biological processes in SCLC. (**A-F**) Bar plots indicating -log(adj.*p* value) of top GO: Biological Processes signatures associated with gene expression modules 1-6 identified in (Figure 6C). Gene ontology analysis was conducted using the g:Profiler web tool. Red bar graphs represent genes enriched in Restored cells after p53 reactivation, while blue bar graphs represent genes enriched in Control cells.

**Fig. S8.**
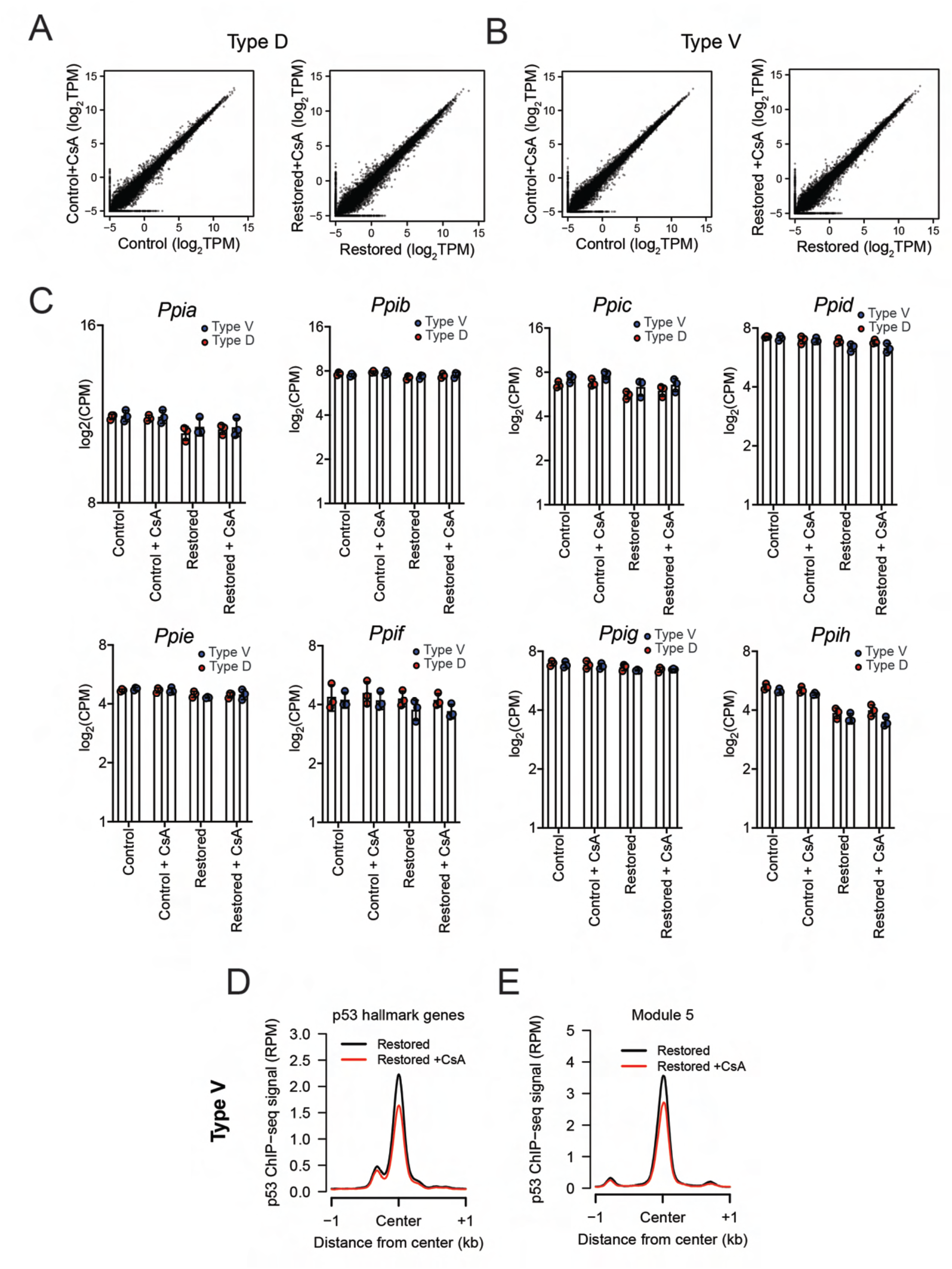
CsA does not influence individual gene transcription or p53 binding. (**A,B**) Dot plot analysis of gene expression 72hrs after CsA treatment in Control and Restored Type D (A) and Type V (B) cells. Mean expression from n=3 cell lines (log_2_ TPM) per represented. (**C**) Quantification of log_2_ counts per million (CPM) of cyclophilin family member transcripts in Control, Restored, Restored + CsA Type D (n=3) and Type V (n=3) cells. Error bars represent mean ± s.d. (**D,E)** Comparison of p53 ChIP-seq signal between Restored and Restored + CsA Type V (n=3) cells for p53 hallmark genes (D) and Module 5 genes (E). Data are centered on p53-bound peaks across a ±1kb window.

